# Sex effects on gene expression across the human cerebral cortex at single cell resolution

**DOI:** 10.1101/2025.06.30.661781

**Authors:** Alex R. DeCasien, Pavan Auluck, Siyuan Liu, Stefano R. Marenco, Mark Cookson, Armin Raznahan

## Abstract

Sex differences in brain-related health outcomes may be a consequence of differences in gene expression. However, most current knowledge relies on studies of bulk tissue or isolated brain regions. Here, we present a large-scale single-cell analysis of transcriptomic sex differences in the human brain, using 169 samples from 15 females and 15 males across six cortical regions, selected based on in vivo neuroimaging measures of sex-biased volume. We find that sex effects on gene expression are highly patterned across cortical regions, cell types, and genes. They are most pronounced in: i) multiple cell types in the fusiform cortex (linked to male-biased volume and sex-biased behaviors); ii) oligodendrocytes, astrocytes, and excitatory neurons across regions; and iii) a subset of sex chromosome and autosomal genes. Over 3,000 unique genes exhibit sex-biased expression, with 133 genes (119 autosomal) showing consistent sex differences across all region x cell type combinations. Sex chromosome genes show the largest sex differences in expression, driven by conserved X-Y gametologs, cell-type-specific biases in certain X- and Y-linked genes, and escape from X-inactivation – with the list of known escapees substantially expanded through our single-cell allele-specific expression analysis. Broader effects of sex on autosomal expression are captured in 13 core signatures with varying cell type vs. region specificity. These signatures are: i) shaped by regional differences in metabolism and laminar architecture; ii) enriched for diverse cellular compartments and biological processes; iii) regulated by sex steroids and X-linked transcription factors; and iv) linked to sex-specific genetic risk factors in sex-biased neuropsychiatric and neurodegenerative diseases. This study substantially advances the breadth, depth, and granularity of knowledge on sex differences in the human brain, and provides a new open data resource to support future research.

## Introduction

Males and females (see Note 1) differ in the prevalence, presentation, and progression of multiple brain disorders across the lifespan. For example, males are more often diagnosed with early-onset neurodevelopmental conditions (e.g., ASD, ADHD), Parkinson’s disease, and ALS [1,2], while females are more often affected by migraine, Alzheimer’s disease, and other dementias [3,4]. These differences likely reflect a mix of biological and social factors, but their cross-cultural consistency [5,6] and stereotyped developmental timing [7] point to a possible role for predisposing sex differences in brain organization. A key challenge in exploring this possibility is the brain’s vast complexity, which creates a large and only partially explored search space for identifying disease-relevant sex differences.

To date, most knowledge about sex differences in human brain organization comes from over three decades of in vivo neuroimaging studies [8–11], supplemented more recently by analyses of gene expression in postmortem bulk tissue [12–18]. These approaches have revealed consistent sex differences in both brain structure and molecular profiles, but they lack cellular resolution. Advances in single-nucleus RNA sequencing (snRNA-seq) have highlighted that many features of brain biology – in both health and disease – are cell type-specific [19–24]. However, this technology has not yet been widely applied to investigate sex differences across multiple brain regions at scale [19–28].

Here, we analyze cell type-specific sex differences in gene expression using a new snRNA-seq dataset spanning 169 samples from six cortical regions in 30 adult individuals (15 females, 15 males) (Figures 1A, 1B; Table S1). To bridge molecular data and neuroimaging, we target six regions chosen for their patterns of reproducible sex differences in cortical grey matter volume (GMV) identified from in vivo structural magnetic resonance imaging (sMRI) in two independent datasets encompassing a total of 2,096 individuals (1,048M/1,048F) (Methods; Figure 1A, Figure S1) [9,29]. Our study addresses several key questions about molecular sex differences in the human brain at single-cell resolution. First, we examine sex differences in mean cell type proportions and gene expression across regions and cell types and test their associations with sex-biased GMV. To capture different dimensions of sex-biased expression, we apply multiple analytic approaches, including: i) variance partitioning [20]; ii) counts of differentially expressed genes [27,28]; and iii) comparisons of effect size magnitude [30]. Second, given the categorical sex differences in X and Y chromosome dosage between XY males and XX females, we provide an unprecedentedly detailed map of sex chromosome gene expression across all major cortical cell types. This analysis reveals a core set of genes showing ubiquitous sex-biased expression, along with cell-type-specific patterns of Y-linked expression and escape from X chromosome inactivation (XCI) – an epigenetic process, regulated by the X-linked lncRNA *XIST*, that randomly silences one X chromosome in each female cell [31]. Third, we provide a systematic survey of sex differences in autosomal gene expression, resolving how these differences vary by cell type and region. Finally, we annotate sex-biased autosomal gene expression for potential regulation by sex chromosomes and sex steroid hormones, and evaluate their contributions to sex-biased genetic risk for neuropsychiatric disorders.

**Figure 1.**
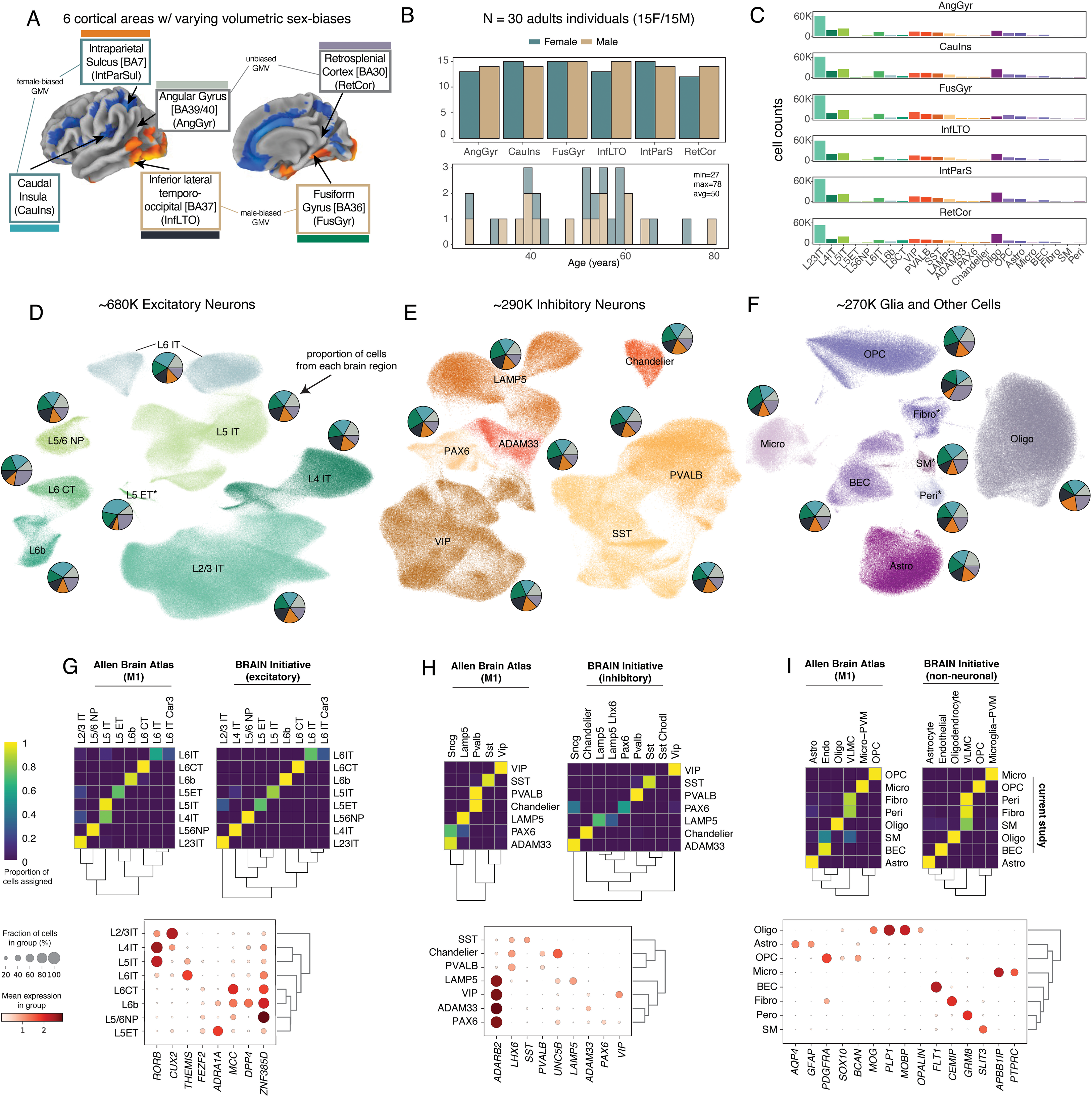
Dataset overview. A) Six brain regions sampled in the current study (right hemisphere shown, see Fig. S1B) [29]. On brains, colors indicate the direction and magnitude of sex-biased grey matter volume (GMV) (blue = female-biased, yellow/orange = male-biased). Colors of the boxes around region names also indicate the direction of GMV sex-bias (male = brown; female = teal, unbiased = grey). Line/block colors represent the different brain regions (AngGyr = gray; CauIns = blue, FusGyr = green; InfLTO = black; IntParSul = orange; RetCor = purple). B) Number of male and female individuals included in each region (top) and age distribution of individuals (bottom). C) Number of cells obtained from each subclass (ordered from most to least within each cell class – excitatory neurons, inhibitory neurons, other). Colors indicate cell subclass. Excitatory neurons are named according to their cortical layer (e.g., L23 = layer 2/3) and projection (IT = intertelencephalic neurons; ET = extratelencephalic; NP = near projecting; CT = corticothalamic neurons). Inhibitory neurons are named according to their marker genes (e.g., PVALB, SST, LAMP5) or morphological types (Chandelier). Glial cell abbreviations are as follows: Oligo = oligodendrocytes; OPC = oligodendrocyte progenitor cells; Astro = astrocytes; Micro = microglia; BEC = brain endothelial cells; Fibro = fibroblasts; SM = smooth muscle cells; Peri = pericytes. D-F) UMAP plots of excitatory neurons (D), inhibitory neurons (E), and glia and other cells (F). Colors indicate cell subclass (see Figure 1C). Pie charts represent the breakdown of each subclass by region (region colors from Figure 1A). Asterisks (*) indicate cell types that were not analyzed for differential expression (too few samples with >30 cells, see Methods). G-I) Top panels: Proportion of cells in each excitatory neuron (G), inhibitory neuron (H), or other (I) subclass (rows) assigned to each reference cell subclass (columns) using Azimuth. Brighter yellow = higher proportions, darker blue = lower proportions (see legend). References include Allen Brain Atlas (M1 = primary motor cortex) and the BRAIN Initiative. Bottom panels: Dot plots of marker gene expression (columns) for each excitatory neuron (G), inhibitory neuron (H), or other (I) subclass (rows). Larger dots = higher fraction of cells in a group express a given gene (see legend). Darker red = higher mean expression in the group (see legend).

The dataset and analyses presented here offer a reference map of transcriptomic sex differences across human cortical cell types and regions with unprecedented detail, substantially advancing our understanding of sex as a neurobiological variable and endorsing its inclusion in future basic and clinical neuroscience research.

## Results

### Dataset overview

The 169 samples used for snRNA-seq analysis were dissected from six cortical regions selected from 30 donors (15F/15M) (Table S1) based on reproducible sex differences in gray matter volume (GMV) across two large in vivo sMRI datasets [9]: two regions show female-biased GMV caudal insula (CauIns) and intraparietal sulcus / BA7 (IntParSul); two show male-biased GMV inferior lateral temporo-occipital cortex (InfLTO) and fusiform gyrus (FusGyr); and two show no significant sex differences in GMV – angular gyrus (AngGyr) and retrosplenial cortex (RetCor) (Figures 1A, S1; Methods; Table S1). We used alevin-fry [32] for snRNA-seq read mapping and gene expression quantification, as it allows retention of multi-mapping reads [33] and more accurate quantification of sex chromosome genes with high sequence similarity between X and Y chromosomes [34] (Tables S2-S4). To minimize contamination in cell type identification and differential expression [35], we removed ambient RNA and called cells using CellBender [36] (Table S4). After sample and cell-level quality control (Methods), our dataset included approximately 1.2 million nuclei across three major cell classes: ∼680k excitatory neurons, ∼290k inhibitory neurons, and ∼270k glial and other cells (Figures 1C-F). Across the three cell classes, we identified N=24 subclasses (Figures 1C-I, S2) and N=86 supertypes (Figure S2) using reference mapping in Azimuth [37–39] (Figures 1G-I, S2) (Methods). Due to the overall sparsity of cells within supertypes (Figure S2) and in line with recent work [20], we focused our analysis of sex differences in gene expression on the 19 subclasses with sufficient cell counts across individuals (Methods).

### Cortical cell type proportions do not exhibit prominent sex differences

We first tested for sex differences in cell type proportions, both within and across brain regions, using a modeling approach that accounts for technical variation and relevant covariates [40] (Methods). Consistent with recent snRNA-seq studies of (primarily) the frontal cortex [20,22] and the largest available histological studies [41,42], we found no evidence of sex differences in the average proportions of any cell type, including neurons versus glia, either across or within regions (all p_adj_>0.05) (Figure 2A; Tables S4-S5). Given our sMRI-guided selection of brain regions, this finding suggests that sex differences in relative GMV may be driven by differences in cellular morphology rather than cellular composition.

**Figure 2.**
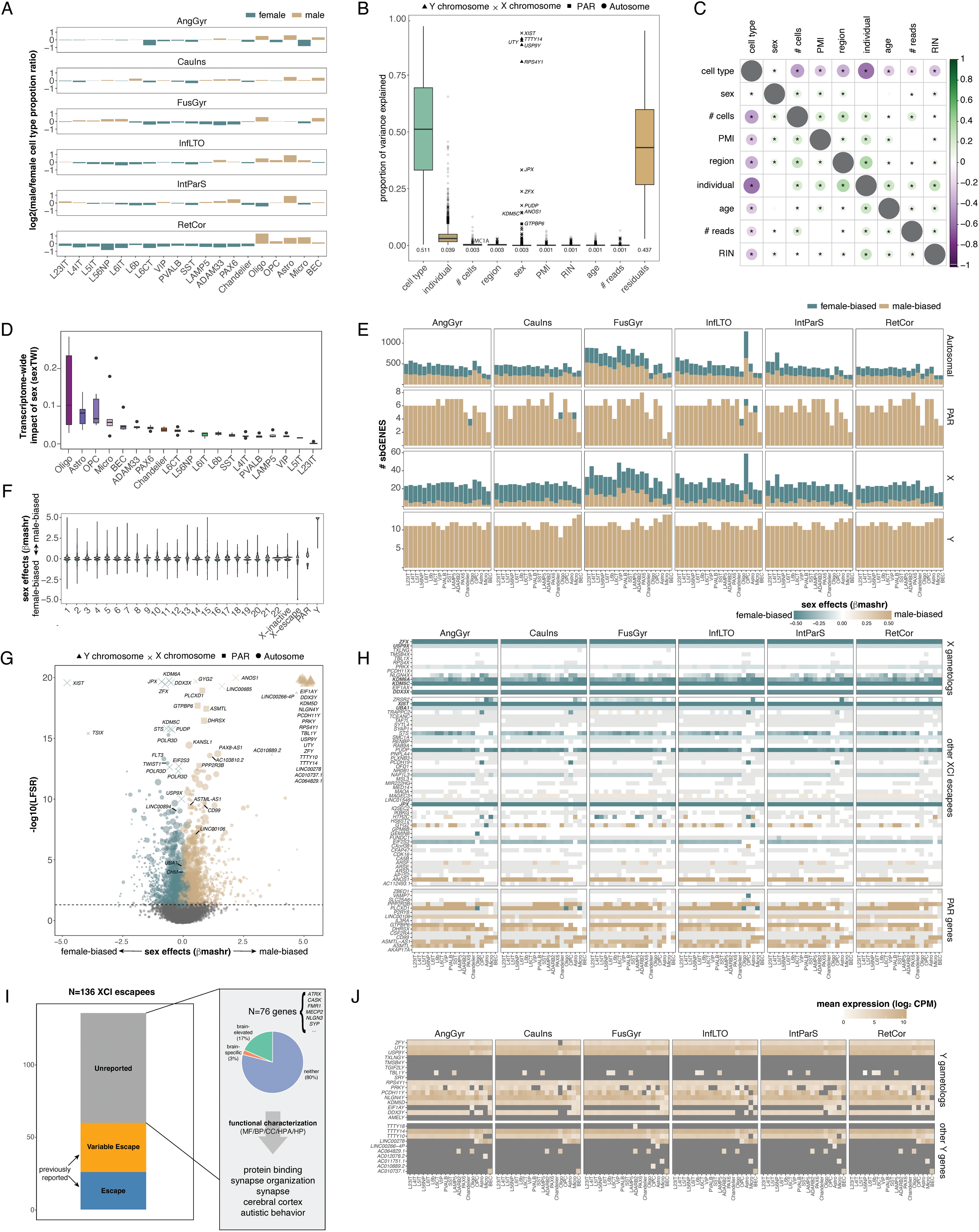
Sex differences in mean expression. A) Ratio of male to female cell type proportions for each region x subclass (log2 transformed ratios; positive = male-biased; negative = female-biased). Color indicates direction of bias (as in Figure 1). None of these differences are significant. B) Variance partitioning plot showing the proportion of gene expression variance explained by each biological and technical factor. ∼4k genes expressed in all samples and subclasses were included. Mean values for each factor are shown below each boxplot. Outlier shapes indicate chromosomal location of the gene (see legend). Examples of genes for which sex explains a high proportion of variance are shown. C) Correlation plot of variance partitioning results. Size of dots indicates strength of correlation. Darker red = more negative. Darker blue = more positive. Asterisks (*) indicate p_adj_ < 0.05. D) Transcriptome wide impact of sex (sexTWI) within each cell subcl;ass (each boxplot contains 6 region-specific sexTWI values). E) Barplots showing counts of sex-biased genes (sbGENES; LFSR < 0.05) grouped by chromosome location (autosomes, pseudoautosomal regions, X chromosome, Y chromosome) and direction of bias (see legend). F) Magnitude of sex effects (β_mashr_) for sbGENES (LFSR < 0.05) across chromosomes and sex chromosome gene subsets (X chromosome genes are separated into those previously reported to be silenced or to escape XCI). Positive = male-biased. Negative = female-biased. G) Volcano plot showing the highest -log10 LFSR value for each gene (across region x subclass combinations) and its corresponding sex effect (β_mashr_). Size of dots corresponds to the number of region x cell type combinations with LFSR < 0.05. Color indicates direction of bias (see legend). Shape indicate chromosomal location of the gene (see legend). H) Sex effects (β_mashr_) for genes belonging to different X chromosome gene sets, including: i) X gametologs; ii) non-gametolog XCI escapees; and iii) PAR genes. Bolded genes are consistently female-biased in all 144 region x subclass conditions. I) Summary of detected XCI escape genes from biallelic expressions analysis. Genes previously reported to escape in [50,51]. Examples of the N=76 unreported N=76 genes are listed, and top enrichment results from g:Profiler are summarized (MF = molecular function; BP = biological process; CC = cellular compartment; HPA = human protein atlas; HP = human phenotype). J) Y chromosome gene expression within each region x subclass combination. Note: *AC010889.2* is an isoform of *KDM5D*; *AC010737.1* is *lnc-TGIF2LY-1*.

### Sex explains a small but distinct proportion of overall gene expression variance

We first assessed the impact of sex on single-cell gene expression in relation to other biological factors – namely cell type, brain region, and age – using a variance partitioning approach. This analysis focused on 4,313 genes expressed across all cell types in each of the 169 samples (Methods, Table S6). As expected, the largest source of variance was cell type (mean=51.1%), followed by individual-level effects (mean=3.9%). In contrast, sex, brain region, and age each contributed a relatively small fraction of the total variance across all genes (sex: mean=0.3%; region: 0.3%; age: 0.1%) (Figure 2B; Table S6). However, consistent with karyotypic differences between XX females and XY males, sex explained a much larger proportion of the variance for sex chromosome genes compared to autosomal genes (Y chromosome: mean=87.6%; X chromosome: 2.0%; pseudoautosomal region [PAR]: 3.3%; autosomes: 0.1%) (Figure 2B; Table S6) [20]. When we repeated the variance partitioning within individual cell types and brain regions, sex accounted for a substantially higher proportion of variance (mean=5.0%, range=2.8-9.6%), with the largest effects observed in glial populations and the smallest in excitatory neurons (Figure S3; Table S7).

To compare the effects of sex on gene expression variance with those of other biological factors, we correlated the variance explained by each pair of variables across genes (Figure 2C; Table S8). Genes for which sex explained more variance tended to show greater region-specific expression and less cell type-specific expression (sex vs. region: Spearman’s ρ=0.12, p_adj_=1.11e-14; sex vs. cell type: ρ=-0.07, p_adj_=2.30e-5), suggesting that sex differences are more regionally localized than cell type-specific. We found no correlation between the variance explained by sex and by age (ρ=-0.02, p_adj_=0.18), indicating largely independent effects of these two variables. Finally, we observed a strong negative correlation between the variance explained by cell type and by individual (Figure 2C; Table S8), consistent with the expectation that genes with strong cell type specificity are more consistent across individuals.

### The distribution of sex differences in gene expression across cell types and brain regions

We next quantified sex differences in gene expression within each brain region and cell type. Using linear models that controlled for age and technical covariates (Methods), we estimated the average sex effects on gene expression for each region x cell type combination (“condition”) and applied multivariate adaptive shrinkage (MASH) to increase statistical power (Figure S4), improve precision, and compute local false sign rates (LFSRs) [43,44] (Table S9), in line with previous work [12,45]. LFSRs reflect the confidence in the direction of an effect, while accounting for multiple comparisons across tests [44]. MASH identified both condition-specific effects (i.e., individual region x cell subclass combinations; e.g., ILTO OPC-specific, FusGyr Micro-specific) and shared effects across multiple conditions (e.g., global effects shared across all conditions, cross-region glia effects, and cross-cell type FusGyr effects) (Figure S4).

Consistent with our variance partitioning results (Figure S3; Table S6), glia exhibited the largest average absolute sex effects across the transcriptome (|β_mashr_| regardless of LFSR) (ANOVA: region F=114.4; cell type F=96.5; both p<2.2e-16) (Figure S5; Table S9) (pairwise Tukey HSD results in Table S10). Glia also showed the strongest global transcriptome-wide impact of sex (sexTWI) (Figures 2D, S6; Table S11) (ANOVA: p = 1.29e-11; Tukey’s HSD: p_adj_<0.05 for glia > excitatory and inhibitory neurons), a summary measure of the overall impact of sex on gene expression in each condition (Methods) [46]. Brain regions with male-biased GMV (FusGyr, InfLTO) tended to show the highest sexTWI across cell subclasses, followed by regions with non-biased GMV and then those with female-biased GMV (Figures S6, S7A; Table S12). Consistent with this pattern, sex-biased GMV (positive = male-biased; negative = female-biased) was positively correlated with sexTWI, though the association reached nominal significance (p < 0.05) in only 2 of 19 cell subclasses (Figure S7A; Table S12). Subsampling analyses (Methods) produced similar rankings of region x cell type combinations by sexTWI (correlation (ρ) between original ranks and median rank across 20 iterations = 0.328, p = 3.632e-4), although subsampling-derived sexTWI tended to be lower for oligodendrocytes and microglia in some regions (Figure S6).

We identified thousands of sex-biased genes (N=3,382 sbGENES; LFSR<0.05) across our dataset (Figure 2E; Table S9). 133 sbGENES were sex-biased in all 114 conditions, while more genes were biased in individual regions (N=1,455) or in individual cell types (N=1,337) (Tables S9, S13). These 133 genes appear to represent ubiquitous sex differences in gene expression in the human brain, with 14 located on the sex chromosomes and the majority being autosomal (Table S13). Both ubiquitously female-biased (N=61) and male-biased (N=58) autosomal genes were linked to neurodevelopmental abnormalities, macromolecule metabolic processes, and gene expression regulation (g:Profiler GO:BP, g:SCS<0.1) (Table S14; Methods). Interestingly, ubiquitously male-biased genes were also enriched for testis expression (g:Profiler HPA, g:SCS=3.43E-11) (Table S14), suggesting pleiotropic targets of sex-biased biology shared across tissues. Variability in the proportion of expressed genes that were sbGENES was greater across regions than cell types (ANOVA: region: F=14.85, p=1.3e-10; cell type: F=3.25, p=1.07e-4). The FusGyr and InfLTO – the two regions with male-biased GMV (Figure 1A) – had the highest proportions of sbGENES (Tables S9, S15) (Figures 2E, S8A). Consistent with this pattern, sex-biased GMV (positive = male-biased; negative = female-biased) was positively correlated with the proportion of sbGENES, though the association reached nominal significance (p < 0.05) in only 1 of 19 cell subclasses (Figure S7B; Table S12). Among cell types, L23IT, L5IT neurons, and OPCs showed the greatest proportions of sbGENES (Tables S9, S15) (Figures 2E, S8A). Subsampling analyses (Methods) detected fewer sbGENES (due to loss of power) but produced similar ranks of cell type x region combinations according to the number of sbGENES (correlation (ρ) between original ranks and median rank across 5 iterations = 0.753, p < 2.2e-16; subsampling-derived sbGENE counts tended to be lower for OPCs and astrocytes in some regions) (Figure S8B).

Thus, while variance partitioning (Figure S3; Table S6), absolute effect sizes (Figure S5; Table S9), and TWI measures (Figure 2D; Table S11) indicated that glial cells are most affected by sex, sbGENE counts from the MASH analysis revealed strong sex effects on neuronal gene expression across multiple regions, as well as on OPCs in the InfLTO (Figures 2E, S8A; Table S9). This likely reflects two factors: i) neuronal subclasses are more numerous and have more similar expression profiles, increasing statistical power in MASH meta-analysis; and ii) glial cells show greater interindividual variability (Figure S8D), leading to less precise pre-MASH estimates (Figure S8E). Together, these findings illustrate the complementary strengths of different analytic approaches in capturing distinct aspects of transcriptomic sex differences, and collectively point to specific cell types and regions that may be especially sensitive to sex.

### Sex chromosome genes exhibit the largest sex differences in expression

Amongst sbGENES, those on the sex chromosome exhibited the largest sex differences in expression across brain regions and cell types (Figures 2F-H; Table S9), consistent with our variance partitioning results (Figure 2B). Genes in the pseudoautosomal regions (PARs) of the X and Y chromosomes were predominantly male-biased (Figures 2E–H; Table S9), consistent with previous reports from bulk RNAseq studies [12]. This sex bias may reflect incomplete escape from XCI for PAR genes on inactivated X-chromosomes in females in comparison with the potential for full expression of these genes from both the X and Y copy in males [47]. *PLCXD1* – the most telomeric PAR gene and therefore perhaps least exposed to spreading of XCI on the inactive X chromosome (X_i_) – deviated from this pattern in some cell types, showing female-biased expression in oligodendrocytes and microglia (Figure 2H). Outside the PAR, we observed variable sex differences in expression across different X-linked genes (Figure 2H; Table S9). Ten X-chromosome genes displayed ubiquitous female-biased expression across all region x cell type combinations (*CHM, DDX3X, JPX, KDM5C, KDM6A, UBA1, ZFX, USP9X, XIST, LINC00894*; *CHM* and *UBA1* show relatively small female-biases) (Figure 2G; Table S9). These genes included effectors of X chromosome inactivation (XCI) (*XIST*, *JPX*) and highly constrained X gametologs that escape XCI (*DDX3X*, *KDM6A*, *KDM5C*, *ZFX*, *USP9X*). The latter genes have retained non-recombining copies on the Y chromosome and show extreme expression sensitivity to variations in sex chromosome dosage [48,49].

For X-linked genes that did not show ubiquitous female-biases in expression, observed variation in the degree of female-biased expression partly cohered with past reports of XCI status [50,51] (Figure 2I). Notably however, some X gametologs previously reported to escape XCI showed no sex bias (*TXLNG*, *RPS4X*, *EIF1AX*), potentially reflecting brain-specific regulation [51]. Other gametologs reported to exhibit variable XCI escape [34,52] showed inconsistent (*NLGN4X* and *PRKX*) or absent (*TBL1X*, *TMSB4X*, *PCDH11X)* female-biases in gene expression across region x cell type combinations. Non-gametolog XCI escapees exhibited cell type-specific female biases in expression (e.g., *STS*, *EIF2S3*, and *NAP1L3* in non-glial cells; *ZRSR2* in glia). We also observed restricted female-biased expression of *TRAPPC2* and *PCDH19* in OPCs. A few non-PAR X-linked genes with variable or disputed XCI escape status showed male-biased expression (e.g., *GYG2*, *ANOS1*, *ARSF*, *MAGEC3*) [50,51,53,54] (Figure 2G; Table S9).

Overall, these patterns of sex-biased X chromosome gene expression align with prior studies of sex-biased expression and XCI escape [50,51,53,54], but also highlight the potential for context-dependent regulation across cell types and brain regions.

### Expanding the inventory of brain-expressed genes that escape X chromosome inactivation

While the patterns of sex-biased X chromosome gene expression observed in this study generally align with previous classifications of genes based on their XCI status, the latter are largely based on bulk studies of tissues besides the brain [47,50,51,55,56]. This gap in knowledge limits our understanding of how patterns of XCI may differ between the brain and other tissues, and constrains mechanistic insights into sex differences in X-linked gene expression in the brain. For example, female-biased expression could reflect escape from XCI (XCIe, where a gene is mono-allelicaly expressed in males and bi-allelicaly expressed in females), but it could also arise from XCI-independent differences in mono-allelic expression of a gene in males and females (e.g., upregulation in females from the active X chromosome [X_a_] alone). To address this, we performed allele-specific expression (ASE) analyses (Methods) to conduct the first systematic survey of XCIe across brain cell types. To validate our approach for distinguishing monoallelic versus biallelic expression of sex chromosome genes in females, we confirmed that it accurately classified PAR genes as biallelic and non-PAR X-linked genes as monoallelic in males (Figure S9; Table S16; Methods). Given this validation, we identified XCIe genes as those with evidence of biallelic expression in at least one cell type from our cohort of female donors (Figure S9; Table S16; Methods). We found evidence for XCIe for several well-established XCIe genes (e.g., *ZFX*, *USP9X*, *KDM5C*, *UBA1*), but also for several novel or rarely reported XCIe genes (e.g., *ATRX, CASK, FMR1, MECP2, NLGN3, MAGED1, SYP, TSPAN7, TSPYL2)* (Table S16). These novel XCIe genes tend to be expressed in the brain and are associated with synaptic functions and ASD (p_adj_ < 0.1) (Figure 2I; Table S17). Furthermore, many of these newly discovered XCIe genes did not show female-biased expression in our dataset (e.g., *ATRX, SYP, FMR1*) (Tables S9, S16), illustrating the limitations of using sex-biased expression alone to infer XCI status. The expanded list of XCIe genes presented here (Table S16) is likely to represent a lower bound for XCIe in the brain, which would likely increase with deeper snRNA-seq coverage.

### Y chromosome genes exhibit cell type-specific expression patterns

We found that Y chromosome genes exhibit diverse expression patterns across brain cell types in XY males (Figure 2J; Table S9). Some Y gametologs were consistently expressed across all 114 region x cell type combinations (e.g., *UTY*, *USP9Y*, *RPS4Y1*, *TTTY14*), while others exhibited variable expression levels (e.g., *EIF1AY*, *NLGN4Y*, *PRKY*) (Figure 2J; Table S9). These patterns largely mirrored the female-biased expression observed for their X-linked homologs (see above) (Figure 2H; Table S9). Two Y-linked gametologs showed cell-type specific expression patterns, including *TBL1Y* in specific neuronal subtypes (e.g., L6CT excitatory, LAMP5 inhibitory) and *EIF1AY* in multiple non-neuronal cell types (e.g., Oligo, BEC) (Figure 2J; Table S9). We did not detect expression of *TMSB4Y* or *TXLNGY* (Figure 2J; Table S9) – contrary to previous bulk RNA studies [57] – but this was consistent with expectations since *TMSB4Y* is very lowly expressed [57] and *TXLNGY* is a transcribed unprocessed pseudogene that was filtered out of our reference (Methods).

Y-linked lncRNAs also showed variable expression across cell types, with patterns that were generally consistent across cortical regions. Two multicopy lncRNAs involved in RNA splicing (*TTTY10* and *TTTY14*) were broadly detected across cell types and regions (Figure 2I; Table S9). Although traditionally associated with spermatogenesis – to the point of being named “testes-specific transcripts” – these TTTY lncRNAs have been reported in the brain and other somatic tissues and linked to multiple disease, including cancer [58–60]. Most other Y-linked lncRNAs were more cell-type specific (e.g., *LINC00278* was expressed across glia, *LINC00266-4P/SEPTIN14P23* was expressed astrocytes), consistent with prior findings that lncRNAs tend to exhibit greater cell type specificity than protein-coding genes [61,62].

Together, this survey of Y chromosome gene expression across brain regions and cell types offers a foundation for exploring male-specific genetic contributions to brain biology and highlights targets for future investigation into cell type-specific Y-linked regulatory mechanisms.

### Sex-biased gene expression is coordinated between brain regions and cell types, shaped by laminar architecture, and intersects with diverse biological process and cellular compartments

Although the largest sex differences in gene expression across brain regions and cell types were observed on the sex chromosomes (Figure 2F-H), the majority of sbGENES identified in our analyses were located on autosomes (Figure 2E). We therefore next investigated how autosomal sex effects are patterned across cortical regions and cell types. Correlations of autosomal sex effect estimates (β_mashr_ regardless of LFSR) across all region × cell type combinations revealed that sex-biased gene expression is often highly coordinated between regions and cell types (Figure 3A; Table S18). Clustering this correlation matrix identified 13 distinct signatures of autosomal sex-biased gene expression, each shared by specific sets of regions and/or cell types (Figures 3A, S10; Table S19) (Methods). These included: i) region-specific clusters (e.g., Cluster 3 = neurons and glia in FusGyr); ii) cell type-specific clusters (e.g., Cluster 4 = astrocytes across regions; Clusters 5 and 6 = OPCs and oligodendrocytes across regions; Cluster 7 = BECs across regions); and iii) region x cell type clusters (e.g., Clusters 8, 12, and 13 separate inhibitory vs. excitatory neurons in CauIns and RetCor; Clusters 2 and 9 separate Chandelier cells and lower layer excitatory neurons vs. upper layer excitatory neurons in AngGyr, CauIns, IntParS, and InfLTO) (Figure 3A; Table S19). Microglia did not form a distinct cluster but grouped with other glial cell types (Figure 3A; Table S19). Clusters that included similar cell types (e.g., Clusters 5 and 6 = OPCs and oligodendrocytes across different regions) or overlapping regions (e.g., Clusters 2 and 9 = various neuronal subtypes in AngGyr, CauIns, IntParS, InfLTO) showed the strongest inter-cluster correlations (Figure 3B; Table S19). These findings suggest that the transcriptional programs underlying sex-biased gene expression can show sensitivity to both region and cell subclass.

**Figure 3.**
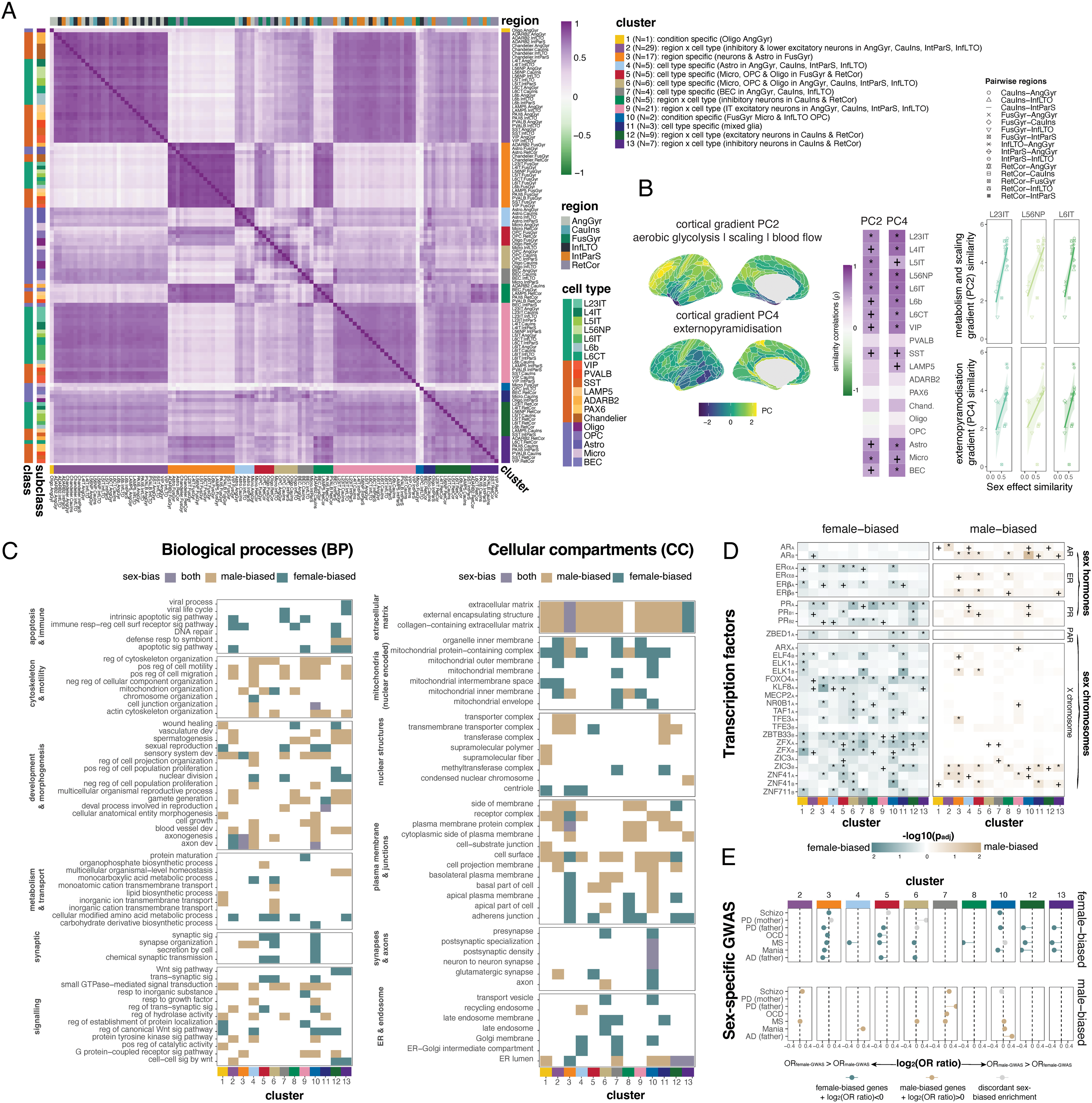
Autosomal sex-biased genes. A) Correlation plot of sex effects (regardless of LFSR) across region x subclass combinations. N=13 clusters were identified via block modelling (Methods). See legends for regions, cell classes, cell subclasses, and cluster descriptions. B) Left: cortical gradient of PC4, which primarily reflects externopyramisation (see Figure S11); Middle: Correlations between PC4 similarity and sex effect similarity within cell subclasses (see legend). Asterisks (*) indicate p_adj_<0.05. Plus signs (+) indicate nominal p<0.05. Right: Scatterplots showing PC4 similarity (y-axis) versus sex effect similarity (x-axis) across region pairs (shapes, see legend) for 10 cell subclasses with significant correlations (p_adj_<0.05 in the middle panel). C) Enrichment results for biological processes (y-axis; left) and cellular compartments (y-axis; right) – organized by function/location (bold groupings on the y-axes) – for each of the 13 clusters from Figure 3A (x-axes). For each cluster and sex bias (male or female), the top 3 terms (highest fold enrichments and p<0.05) were selected, and the results for this union set of terms across clusters were visualized. Colors of tiles represent the sex-biased gene set for which the enrichment was significant (see legend; “both” = male- and female-biased genes in a given cluster were both enriched for that term). D) Enrichments for transcription factors (TFs; y-axis) among female-biased (left) and male-biased (right) genes in each of the 13 clusters from Figure 3A (x-axes). AR = androgen receptor; ER = estrogen receptor; PR = progesterone receptor; PAR = pseudoautosomal region. Subscripts on TFs: A = HOCOMOCO TF dataset; B = Homer TF dataset. Darker colors = stronger enrichment (lower p_adj_ value). Asterisks (*) indicate = p_adj_<0.05; Plus signs (+) indicate nominal p<0.05. Colors show the direction of sex bias. E) Relative enrichment for sex-specific GWAS hits for various sex-biased conditions (y-axes) among female-biased (top) and male-biased (bottom) genes in the 13 clusters from Figure 3A (clusters without significant enrichments are not shown). Here we show the log_2_ ratio of the odds ratios from Fisher’s exact tests for overlap with male-specific GWAS versus female-specific GWAS (OR_male-GWAS_/OR_female-GWAS_) (closest gene to SNP) for each disease and sex-biased gene set combination. This measure represents the relative enrichment for male-vs. female-specific ^GWAS (ratios > 0 = OR^male-GWAS ^> OR^female-GWAS^; ratios < 0 = OR^male-GWAS ^< OR^female-GWAS^). We^ visualized these ratios when either OR_male-GWAS_ or OR_female-GWAS_ was significant (nominal p<0.05). Relative enrichment of sex-specific GWAS signals for various sex-biased conditions (y-axis) among female-biased (top) and male-biased (bottom) genes across the 13 clusters from Figure 3A (only clusters with significant enrichments are shown). We plot the log₂ ratio of odds ratios from Fisher’s exact tests comparing overlap with male-specific versus female-specific GWAS hits (i.e., log₂[OR_male-GWAS_ / OR_female-GWAS_], based on the closest gene to each SNP). Positive values indicate greater enrichment in male-specific GWAS, while negative values indicate greater enrichment in female-specific GWAS. Ratios are shown only when either the male-or female-specific GWAS overlap was nominally significant (p < 0.05).

Some signatures of sex-biased autosomal gene expression appeared to be regionally localized—for example, Cluster 3 was enriched in the FusGyr, while Clusters 12 and 13 were largely restricted to the CauIns and RetCor. This spatial patterning led us to hypothesize that the similarity of sex effects on gene expression between two regions within a given cell type might reflect broader similarities in cortical organization. To test this, we leveraged a recently developed multimodal resource that integrates neuroimaging and postmortem data [63], providing spatially aligned cortical maps of diverse structural and functional features (Figure S11A). These features, which vary along the sensory-to-association cortex axis [63], include: 1) the T1/T2 ratio (a proxy for myelination), 2) externopyramidization (relative soma size in upper vs. lower cortical layers), 3) cortical thickness, 4) allometric scaling (regional areal expansion with brain size),(5) evolutionary expansion (humans vs. macaques), 6) a large-scale transcriptomic gradient, 7) cerebral blood flow, 8) aerobic glycolysis, 9) functional hierarchies (resting-state), and 10) functional specialization (task-based fMRI). Because these features are intercorrelated (Figure S11B), we performed principal component analysis (PCA) to reduce dimensionality. The first five principal components (PCs), each explaining ≥5% of variance, collectively accounted for 85% of total variance (Methods; Figure S11C). For each PC, we computed pairwise similarity among the six cortical regions in our dataset and correlated these matrices with inter-regional similarities in sex effects on gene expression, calculated separately for each cell type (Figure 3A, S11; Methods). PC2 and PC4 – primarily reflecting metabolic features and externopyramidization, respectively (Figure S11D) – showed the most consistent positive associations with sex effect similarity across cell types (PC2: padj < 0.05 for 10 cell types; PC4: padj < 0.05 for 10 cell types) (Figures 3B, S11D–E). Externopyramidization, defined as the ratio of supragranular to infragranular pyramidal neuron soma size, increases with cytoarchitectonic differentiation across the cortex [64,65]. Inter-regional similarity in sex-biased gene expression was not explained by regional similarity in sex-biased GMV (Figure S7C; Table S12). These results suggest that shared laminar architecture and metabolism may underlie regional similarities in sex-biased gene expression, especially among excitatory neurons (Figures 3B, S11D).

We next identified the biological processes and cellular compartments associated with the union set of autosomal sbGENES in each cluster (Table S19) using gene ontology enrichment analysis (Methods’ Tables S20-S21). Here, we focus on the top enriched terms from each cluster (p < 0.05 and top 3 highest fold enrichments) (Figure 3C; Tables S20-S21). sbGENES were broadly involved in immunity, cytoskeletal and synaptic organization, signalling, morphogenesis, and metabolism (Figure 3C), and they were associated with synapses, axons, membranes, endosomes, and nuclear, mitochondrial, and extracellular structures (Figure 3C).

We observed a striking enrichment of extracellular matrix (ECM)-related terms among sbGENES across nearly all 13 clusters, underscoring the extracellular compartment as a common site of sex-biased gene expression across diverse cortical regions and cell types (Figure 3C). ECM terms were typically enriched among male-biased genes, with two notable exceptions: i) Cluster 3 (FusGyr neurons and astrocytes), where both male- and female-biased sbGENES showed ECM enrichment; and ii) Cluster 7 (inhibitory neurons), where enrichment was specific to female-biased genes (Figure 3C; Tables S20-S21). Other top enrichments among sbGENES that recurred across multiple clusters included: i) predominantly male-biased enrichment of cell surface markers, regulation of cytoskeletal organization, sensory system development, and GTPase-mediated signal transduction; and ii) predominantly female-biased enrichment of adherens junction components, mitochondrial protein complexes, regulation of Wnt signaling, and synaptic signaling (Figure 3C; Tables S20-S21).

We also identified cases where the same top biological process or cellular compartment was enriched among both male- and female-biased sbGENES (e.g., axonogenesis and the extracellular matrix in multiple fusiform cell types; Cluster 3) (Figure 3C; Tables S20-S21). These findings suggest that certain aspects of biological organization may be subject to sex-specific regulation or functional compensation.

Most of the top enrichments for sbGENES were specific to individual clusters. For example, Clusters 2 and 9, which captured sex-biased transcriptomic effects in neurons across four brain regions, showed: i) male-biased enrichment of tyrosine and G-protein kinase signaling, blood vessel development, cell motility, glutamatergic synapses, and the nuclear transporter complex; and ii) female-biased enrichment for apoptotic signaling and mitochondrial pathways (Figure 3C; Tables S20-S21). Cluster 3, which reflected sex-biased gene expression in FusGyr neurons and astrocytes, showed: i) male-biased enrichment of synapse organization, G-protein coupled receptor signaling, and mitochondrial organization; and ii) female-biased enrichment of functions related to the endoplasmic reticulum lumen and the apical and basal plasma membranes (Figure 3C; Tables S20-S21).

Taken together, these results reveal the complex landscape of sex-biased autosomal gene expression across diverse cell types and regions of the cerebral cortex. The observed patterns highlight specific biological processes that are enriched for sex-biased expression and may mediate sex-specific responses to genetic and environmental influences on brain function.

### Sex-biased autosomal genes are regulated by sex hormones and sex chromosome transcription factors

Sex differences in autosomal gene expression could potentially reflect downstream impacts of sex differences in gonadal function, sex chromosome dosage, or external environmental factors. To test the first two of these causal hypotheses, we asked if the promoters of autosomal sex-biased genes in each cluster were enriched for elements responsive to steroid hormones and transcription factors (TFs) on the sex chromosomes using two TF datasets [66,67] (Methods). We found that female-biased genes were primarily regulated by progesterones, estrogens, and numerous X chromosome genes (Figure 3D; Table S22) – including *ZFX*, an X gametolog that shows female-biased expression (Figure 2H; Table S9) and escapes XCI in our data (Figure S8; Table S16). In contrast, male-biased genes were primarily regulated by androgens and certain zinc finger proteins on the X chromosome (*ZIC3*, *ZNF41*) (Figure 3D; Table S22). Our observation that female-biased genes were regulated by *ZFX* is consistent with the roles of *ZFX/Y* as transcriptional activators [68]. The motif databases we queried do not include a *ZFY* motif [67]. Despite the male-specific expression of *ZFY* (Figure 2J) and its 97% sequence similarity with *ZFX* in the zinc-finger domains [69], no *ZFX* enrichment was observed among male-biased genes in any of the clusters (all p_adj_>0.05) (Figure 3D; Table S22). Regulation by sex steroids is also consistent with our functional enrichment results (discussed above), including female-biased intracellular estrogen receptor signaling (Clusters 10, 11; e.g., *ESR1*, *CARM1*, *TAF7*) and male-biased hormone metabolism/secretion (Clusters 2-4, 8, 10, e.g., *HSD17B3*, *CRHBP*) (Table S19). These findings suggest that sex-biased gene regulation is not only influenced by sex hormones but also by transcription factors from sex chromosomes.

### Sex-biased autosomal genes are associated with sex-specific common genetic risk for sex-biased diseases

We tested whether sex-biased autosomal genes were enriched for sex-specific common genetic risk associated with several sex-biased neurodevelopmental, psychiatric, and neurodegenerative disorders – including schizophrenia, OCD, schizophrenia, mania, Parkinson’s disease (PD), Alzheimer’s disease (AD), and multiple sclerosis (MS) – using sex-stratified GWAS data from the UK Biobank [70] (see Methods; sex-specific GWAS was not available for ASD). We found that sex-biased genes tended to be more strongly enriched (with a higher odds ratio, OR; M/F OR ratio = log_2_(OR_male-GWAS_/OR_female-GWAS_)) for risk loci identified in the GWAS corresponding to their matched sex (Figure 3E; Table S23), implicating both overlapping and distinct cell subclasses. For example, female-biased genes were significantly enriched (p<0.05) for MS risk genes from the female-specific GWAS in Cluster 3-6, 8, 12, and 13, and these enrichments were stronger than those for the male-specific GWAS (M/F OR ratios < 0) (Figure 3E; Table S23). Stronger enrichment from the male-specific MS GWAS (M/F OR ratio > 0) was only observed in Cluster 10 (Figure 3E; Table S23). Conversely, male-biased genes were enriched for MS risk genes from male-specific GWAS in Cluster 2, 6, 7, and 10, and these enrichments were stronger than those for the female-specific GWAS (M/F OR ratios > 0) (Figure 3E; Table S23). Notably, the common Cluster 6 includes OPCs and oligodendrocytes, which are directly impacted in MS [71] (Figure 3A). Furthermore, Cluster 4–7 and 10 include microglia (Figure 3A), which contribute to neuroinflammation in MS [72]. Several clusters showed enrichment for multiple conditions, which may reflect shared genomic architecture across diseases [73,74]. For instance, female-biased genes in Cluster 3 and 5 were enriched for 6 and 4 disorders, respectively, while male-biased genes in Cluster 7 and 10 were each enriched for 4 disorders (Figure 3E; Table S23). Interestingly, male-biased genes in multiple clusters are enriched for processes implicated in neurodegeneration, including dopaminergic signalling (clusters 1-3, 6-13; e.g., *DRD2*, *LRRK2*) and amyloid fibril formation (clusters 3, 8; e.g., *SERF1A/B*) (Table S19). Together, these results support the idea that sex differences in gene expression may modulate the functional impact of disease-associated variants [13,17,75].

## Discussion

Our study advances the understanding of sex differences in human brain organization by applying snRNAseq to map sex effects on gene expression across multiple cortical regions at single-cell resolution.

First, by targeting cortical regions previously identified through in vivo neuroimaging studies as showing sex-biased differences in cortical volume [9,29], we were able to test for alignment between sex differences at the macroscopic morphological and microscopic molecular scales. We do not detect sex differences in cellular proportions in any of the sampled regions (Figure 2A), suggesting that volumetric sex differences may reflect variation in cellular morphology or intercellular space (Figure 3C) rather than cell type composition. While we cannot rule out that technical variability in snRNAseq or limited sample size may obscure true differences in cellular proportions, our findings are consistent with prior histological studies that also failed to detect sex differences in cell type proportions [41,42]. Among the six cortical regions profiled, the regions with male-biased GMV (fusiform gyrus, inferior lateral temporo-occipital cortex) exhibit the most prominent sex differences in gene expression, while those with female-biased GMV (caudal insula, intraparietal sulcus) exhibit the least (Figure S7; Table S12). This points to the fusiform gyrus as a hotspot for sex differences, since it not only exhibits reproducible sex differences in volume [9,29], but is specialized for face processing – a domain with well-established sex differences [76,77] – and is implicated in autism, a condition showing strongly sex-biased prevalence [78,79]. Given the challenges associated with histological screening of sex differences across the entire human brain, our findings suggest that the fusiform gyrus represents a promising target for future studies of sex-biased neurobiology.

Second, we systematically characterize the impact of sex on gene expression using multiple complementary approaches that allow us to contextualize these effects relative to other sources of variance and examine how they vary across regions, cell types, and genes. For most genes, sex explains a small proportion of expression variance – typically <1% (Figure 2B) – consistent with prior studies [20,80]. However, the impact of sex is markedly higher for sex chromosome genes, consistent with the categorical difference in sex chromosome dosage between XX females and XY males (Figures 2B, 2F). Within these genes, we observe a core set of 14 PAR, X-, and Y-linked genes that show consistent sex differences in mean expression across all regions and cell types (Figure 2H). These genes are enriched for evolutionarily preserved X-Y gametolog pairs with key regulatory functions [81,82], suggesting that they may mediate broader genome-wide effects. Supporting this idea, transcription factor binding sites for ZFX – a ubiquitously female-biased X gametolog – are enriched among sex-biased autosomal genes (Figure 3D). We also identified sex chromosome genes with targeted sex-biased expression in specific cell types (e.g., *PLCXD1* in oligodendrocytes; *HTR2C*, *PCDH19*, and *TRAPPC2* in OPCs; *TBL1Y* in L6CT excitatory and LAMP5 inhibitory neurons) (Figures 2H, 2J), suggesting cell type-specific consequences of sex chromosome dosage. Moreover, by estimating allele-specific expression across a range of brain regions and cell types, we identify previously unrecognized instances of X-chromosome inactivation escape (XCIe) in the human brain. Over 50% of the genes exhibiting biallelic expression in females had not been previously reported to escape XCI in humans, and these newly identified genes are significantly enriched for brain-specific expression, synaptic function, and lack of sex-biased expression. For instance, *SYP* — a presynaptic marker predominantly expressed in the brain [83] — shows evidence of XCIe in our dataset (Table S16) and consistent but non-significant male-biased expression across all region-by-cell type combinations (β_mashr_ < 0 in all region x cell types; all LFSR > 0.05) (Table S9), in line with previous studies [12,51,82]. Notably, XCIe for *SYP* has previously been reported only in mouse brains [84,85] and in a limited number of human-mouse hybrid cell lines [47]. Furthermore, we find evidence that all 10 ubiquitously female-biased X chromosome genes show evidence of XCIe in our dataset (Figure S8; Table S16) – including *CHM* and *LINC00894*, which previously showed inconclusive or inconsistent evidence of inactivation [50,51]. Notably, *LINC00894* plays a role in synaptic function [86] and is directly regulated by estrogen [87], highlighting it as a promising candidate for future studies of brain sex differences. Finally, while *GYG2* generally displays male-biased expression (Figure 2G; Table S9), we observe both female-biased expression and XCIe in astrocytes (Figures 2H, S8; Tables S9, S16). Together, these findings provide a new empirical foundation for understanding how sex chromosomes contribute to sex-biased neurobiology across diverse cell types and cortical regions.

Third, while the largest and most consistent transcriptomic effects of sex localize to sex chromosomes – as commonly reported using classical analytic approaches [12,13,15,16] – our design enables greater power to detect many autosomal sbGENES by leveraging shared patterns across regions and cell types. This approach reveals that sex influences the expression of >3K unique autosomal genes and that 119 of these genes show ubiquitous sex differences across all region-by-cell-type combinations (Table S9), indicating widespread sex-biased regulation of gene regulatory programs and metabolic processes (Table S14). Using dimension reduction on expression fold change profiles across 114 region-by-cell-type combinations, we identify 13 distinct patterns (i.e., clusters) of sex effects on autosomal gene expression. Some clusters are shared across regions within the same cell types (e.g., Clusters 2 and 9), while others span multiple cell types within the same region (e.g., Cluster 3; Figure 3A). In neuronal populations, inter-regional similarity in sex effects is closely tied to externopyramidization (Figures 3B S10) – a histological measure reflecting a region’s position in the cortical processing hierarchy [65]. This observation suggests that regional circuit architecture – particularly the balance of feed-forward and feedback elements – may influence how sex modulates gene expression in neurons. Together, these findings highlight the importance of profiling transcriptomic sex differences across diverse cell types and brain regions, and provide a comprehensive open resource to support broader integration of sex-based analyses in human brain single-cell research.

Fourth, our multi-region single-cell survey reveals that transcriptomic sex biases span a wide array of cellular compartments and biological pathways, including those involved in cellular energetics, signaling, connectivity, and the extracellular matrix (ECM) (Figure 3C; Tables S20-S21). These broad effects suggest potential for both dynamic amplification and compensatory balancing of sex differences across different spatial and temporal scales. The prominent enrichment of ECM-related gene sets is particularly noteworthy, as it may reflect a molecular correlate of the macroscopic sex differences in cortical volume that motivated our snRNAseq sampling strategy. While our observational study does not allow us to establish causality, the enrichment of transcription factor binding motifs for sex-linked genes (e.g., *ZFX*) and hormone receptors among sex-biased genes (Figure 3D; Table S22) strongly implicates both sex chromosome dosage and gonadal hormones in shaping gene expression. Still, only a minority (133/3,382; ∼4%) of sex-biased genes show consistent effects across all conditions (Figure 2G; Table S9), suggesting that sex differences in gene expression are context-dependent and potentially modulated by environmental or experiential factors [9].

Finally, our findings suggest that sex differences in gene expression may contribute to sex-specific vulnerability to brain disorders. While many neuropsychiatric and neurodegenerative conditions share a largely overlapping genetic architecture between sexes [88], our results show that genes with sex-biased expression are disproportionately enriched for sex-specific genetic risk across multiple disorders (Figure 3E; Table S23). These findings raise the possibility that sex differences in gene expression may modulate the magnitude of genetic effects at risk loci, potentially influencing how disease-associated variants manifest in each sex. In turn, such differences could contribute to the reduced portability or predictive power of polygenic risk scores across sexes, and underscore the need to incorporate sex-stratified expression data into efforts to understand, predict, and treat brain-related disorders. Together, these results highlight transcriptional sex differences as a potentially important mechanism linking sex to differential disease susceptibility across a range of brain conditions.

Our findings and inferences should be interpreted in light of several caveats and limitations. First, although we provide an unprecedentedly large and regionally diverse snRNAseq analysis of sex differences in the cellular organization of the human brain, even larger sample sizes will be needed to disentangle how sex interacts with other factors such as age, endocrine status, and underlying genetic and environmental variation. Second, our study focused on cortical regions that show distinct sex-biased volume in sMRI—enabling comparison between macro- and micro-scale sex differences within the same brain compartments in adults. Future studies should extend this approach to other brain areas – including subcortical regions showing reproducible sex differences in volume [9] – and additional life stages to more fully capture the relationship between cellular and systems-level sex differences. Third, while snRNAseq offers high-resolution insights into cell-type-specific transcriptional profiles, its reliance on nuclear RNA limits the detection of mature transcripts and precludes analysis of post-transcriptional modifications. These methodological constraints should be considered when interpreting transcriptional differences, particularly in relation to protein-level and functional outcomes [9].

Notwithstanding these limitations and caveats, our study provides an initial reference map of transcriptomic sex differences across adult human cortical cell types and regions with unprecedented detail, substantially advancing our understanding of sex as a neurobiological variable and endorsing its inclusion in future basic and clinical neuroscience research.

## Methods

### Tissue selection and dissection

We selected six cortical areas that exhibit a consistent pattern of sex-bias in regional gray matter volume (GMV) in two independent large-scale in vivo structural MRI datasets [9,29], including two female-biased areas (intraparietal sulcus – BA7, caudal insula), two male-biased areas (inferior lateral occipito-temporal cortex – BA37, fusiform gyrus – BA36), and two unbiased areas (angular gyrus – BA39/40, retrosplenial cortex – BA30) as illustrated in Figure S1. We sampled these regions from N=30 individuals (15F/15M) (Figure 1A, S1). The initial dataset included 192 samples (30 individuals × 6 regions = 180 samples, plus 12 technical replicates). Thirteen samples were removed (see below), and the remaining 10 replicates were merged with their corresponding primary samples, resulting in a final dataset of 169 samples (192 – 13 – 10 = 169) (Table S1).

Brain slabs were retrieved from −80C long-term storage on dry ice. Slabs were inspected to identify the ROI(s) and photographed. Slabs were transferred to a dry-ice chilled cutting board in a biosafety cabinet and ROIs were dissected using a dental drill. Post-dissection, slabs were photographed again and all tissues were returned to −80C until nuclei isolation.

### Nuclei isolation

Nuclei were isolated using the Sigma Nuclei PURE Prep Nuclei Isolation Kit protocol. Specifically, frozen samples were physically dissociated in a homogenizing buffer using a dounce homogenizer, 20-25 strokes with a narrow pestle followed by 20-25 strokes with a wide pestle. Chilled lysis buffer was added to the resulting nuclear suspensions and incubated for 10 minutes. A 1.8 M sucrose cushion solution was prepared and added to the bottom of ultracentrifuge tubes. Nuclear suspensions were carefully layered on top of the sucrose cushion and centrifuged for 45 min at 30,000xg at 4C. The supernatant was aspirated and nuclei pellets were resuspended in suspension buffer and washed twice with centrifugation at 500g for 5 minutes at 4C before final resuspension in 110uL buffer.

For nuclei counting, a 10-microliter aliquot of each nuclear suspension was stained 1:1 with 0.4% Trypan Blue. Nuclei were counted using a hemocytometer under a bright-field microscope. Concentrations ranged from 1-1.5 million nuclei per mL. A final aliquot of 50,000 nuclei in one mL of storage buffer was taken for the 10x Genomics nuclei capture.

### Library preparation and sequencing

Gel Bead-in-Emulsions were prepared by loading up to 10,000 nuclei per sample onto the Chromium Chip G (10x Genomics cat#1000073) and run using the Chromium Controller (10x Genomics Version 5.0). cDNA and DNA libraries were generated with Chromium Single Cell 3’ GEM, Library and Gel Bead Kit V3.1 (10x Genomics). cDNA and DNA libraries quality and quantity were assessed by the Qubit and Tapestation (Agilent). DNA Libraries were sequenced using the NextSeq 500/550 High Output Kit v2.5 (Illumina) on an Illumina NextSeq 550 using the 150-cycle High Output kit with either: i) 28 bp read 1 and 91 bp read 2; ii) 101 bp read 1 and 2; or iii) 151 bp read 1 and 2. We translated BCL files to FASTQ files using Illumina *bcl2fastq* v2.20.

### Read mapping and expression quantification

We used Alevin-fry [32] for read mapping and gene expression quantification, and CellBender [36] for cell-calling and correction for ambient RNA contamination.

We first used alevin-fry [32] to map reads and quantify gene expression, since this method allows us to retain multi-mapping reads, which allows Alevin, relative to other approaches (e.g., Cell Ranger), to output a higher number of mapped reads for genes with low sequence uniqueness [33]. This is particularly important for the quantification of sex chromosome genes, some of which have high sequence similarity between homologous members on the X and Y chromosomes [34]. First, we created sex-specific “splici” (spliced+intron) reference indices (masking the Y chromosome for females) using the *make_splici_txome* function from the R package *roe* and the *index* function in *salmon* [32,89]. These references were built following the Cell Ranger human reference (version 2020-A) which uses GRCh38 (GENCODE v32/Ensembl 98) and only includes protein coding genes, lncRNA, and Ig/TcR genes [90]. Then, for each sample, we: 1) mapped the read data (to the appropriate sex-specific index) by running *alevin* in sketch mode with structural constraints (using the *alevin* function in *salmon*); 2) generated a permit list of cells using all possible cell barcodes (using the *generate-permit-list* function from *alevin-fry*; the list of barcodes was obtained from https://teichlab.github.io/scg_lib_structs/data/3M-february-2018.txt.gz); 3) collated all files (using the *collate* function from *alevin-fry*); and 4) quantified UMIs per gene and cell (using the *quant* function from *alevin-fry*) [32,89]. During quantification, we used a strategy similar to the one adopted in Cell Ranger (resolution = cr-like-em), except that it: 1) does not first collapse 1-edit-distance UMIs; and 2) does not discard multi mapping reads, but treats the genes as an equivalence class and determines the counts for each gene via an expectation maximization (EM) algorithm. Since we used “splici” reference indices, this quantification mode separately attributes UMIs to spliced, unspliced (intronic) gene sequence, or ambiguous (not able to be confidently assigned to the spliced or unspliced category). We then created the gene-by-cell expression matrices (per sample) including all UMIs using the *load_fry* function in the R package *fishpond* (which_counts = c(’S’, ‘U’, ‘A’)). Finally, we generated QC reports for each sample using the *alevinFryQCReport* function in the R package *alevinQC* [91] (Table S3).

For each sample, the unfiltered gene-by-cell expression matrix was rounded to the nearest integer (since CellBender expects counts) and then converted from SingleCellExperiment objects to H5Seurat objects (*CreateSeuratObject* function in *Seurat*; *SaveH5Seurat* function in *SeuratDisk*) and then to h5ad files (*Convert* function in *SeuratDisk*). These files were used as input for the *remove-background* function in *cellBender* [36], which was run on GPU with the following parameters (--cuda; --epochs 200 --learning-rate 0.000025 --fpr 0). The number of epochs were increased and the learning rate was decreased after iteratively inspecting the CellBender reports (which showed issues with learning curve convergence). The false positive rate (FPR) was set to 0 following recommendations from the package’s authors, who suggest that in cohort settings where differential expression will be performed downstream, a false positive rate of 0 will help to avoid overcorrection beyond the expected noise budget [36]. For each sample, CellBender identified cells (vs. empty drops which have a posterior cell probability <= 50%) and adjusted expression levels of cells for ambient RNA contamination. This produced a filtered, adjusted gene-by-cell expression matrix for each sample.

### Genotyping and sample ID confirmation

To confirm sample identity, we genotyped a Chromosome 1 from each sample. We extracted and indexed Chromosome 1 from each sample bam file using *samtools* [92] and then calculated allele frequencies and genotype likelihoods using *angsd* [93]. Using these outputs, we obtained estimates of pairwise relatedness between all pairs of samples using *ngsRelate* [94]. We inspected all pairwise relatedness values to confirm sample identities.

### Initial sample- and cell-level QC

We examined all Cell Ranger (v5.0.1) [90] web summaries to identify low quality samples (Table S2). N=7 samples were removed because the program flagged errors (i.e., low nuclei count, low fraction (<70%) of reads in cells) and/or produced a flat barcode rank plot (i.e., barcodes ranked by UMI, which differentiates cells from background when they exhibit a cliff and a knee) with little to no visible clustering. Two additional samples were removed downstream (see below) due to: i) extremely high mitochondrial content across cells; or ii) lack of diverse cell type capture.

For each sample, we removed low quality cells and doublets. Specifically, we converted each filtered CellBender h5 file to a Seurat object (*Read_CellBender_h5_Mat* function in the R package *scCustomize*; *CreateSeuratObject* function in the R package *Seurat*) [95,96] and converted Ensembl gene IDs to gene names (using the features.tsv.gz files generated by Cell Ranger). We removed cells with >5% mitochondrial RNA (following ambient RNA correction). We next normalized expression, clustered cells (resolution=1.2), estimated the expected doublet rate (using the formula: exp_rate = (0.0527 + 0.0008 * # nuclei)/100), and performed doublet detection and removal using *scDblFinder* [97]. Additional QC and cell removal occurred during integration, clustering, and annotation (see below).

Sequencing quality metrics for the samples included in this analysis indicated high data integrity. According to CellRanger, the mean number of reads per nucleus was 30,639, with a median of 3,108 genes and 7,284 UMI counts per nucleus (Table S2). Complementary results from AlevinQC showed an average of 331.5 million reads per sample, with a mapping rate of 81.6% and an estimated mean of 7,508 nuclei per sample (Table S3).

### Integration, clustering, and annotation

We annotated cells at three levels following Johansen and colleagues [20] including cell: i) “class” (excitatory neurons, inhibitory neurons, non-neuronal); ii) “subclass” (e.g., L4-IT, PVALB, oligodendrocytes); and iii) “supertype” (e.g., oligodendrocyte 1, oligodendrocyte 2).

We first merged all sample Seurat objects into one AnnData object (*SaveH5Seurat* and *Convert* functions in R package *Seurat*; *read_h5ad* and *concatenate* functions in python package *scanpy*) [95,98]. We then used *scanpy* to normalize each cell by total counts over all genes (sc.pp.normalize_total), logarithmized the data matrix (sc.pp.log1p), identified highly variable genes (sc.pp.highly_variable_genes), subsetted the data to only include highly variable genes (flavor=’seurat’ [default]), scaled the data to unit variance and zero mean (sc.pp.scale), and performed principal components analysis (sc.tl.pca). We used *scanpy.external* to integrate the data using the Harmony algorithm (sce.pp.harmony_integrate) [99]. We then clustered the data (resolution=1), annotated cell classes (i.e., highest level annotation), and removed low quality clusters (with high mitochondrial content, high doublet scores, low intronic rate, and/or weak/mixed cell type markers). Specifically, we used *scanpy* to compute a neighborhood graph of observations (sc.pp.neighbors; n_neighbors=15, n_pcs=50), performed UMAP (sc.tl.umap), clustered cells using the Leiden algorithm (sc.tl.leiden; resolution=1), and inspected each cluster according to known cell type markers, UMI counts, and mitochondrial content. Cell subclass markers included: oligodendrocytes: *PLP1, OLIG1, OLIG2, MOBP; OPCs: SOX10, CSPG4, PDGFRA*; microglia: *CD74, APBB1IP*; macrophages: *MRC1*; T cells: *PTPRC*; astrocytes: *GFAP, ALDH1L1, AQP4*; neurons: *GRIN1, GRIN2B*; excitatory neurons: *NEUROD6, NRGN, SLC17A6, SLC17A7*, *SATB2*; inhibitory neurons: *GAD1, GAD2, CALB2, PVALB, VIP, SST, LAMP5*; pericytes: *MCAM*; endothelial: *FLT1*; fibroblasts: *DCN* (Figure 1S, S2).

Next, we re-integrated samples within each of the three cell classes (i.e., excitatory neurons, inhibitory neurons, non-neuronal) and performed two additional rounds of clustering, annotation, low quality cluster removal, and re-integration. Once clean clusters were obtained for each cell subclass, we aimed to identify low-level “supertypes”. To define supertypes within each cell class, we first excluded individuals with <2 SDs of the mean number of cells for that cell class and then clustered the cells at increasing resolutions until: 1) all expected subclasses were recovered (Figure S2; Table S4); and then 2) stopped when all individuals were not included in all clusters (excitatory neurons: resolution = 1.8; inhibitory neurons: resolution = 1.0). For nonneuronal cells (which had high intersample heterogeneity in supertype capture), we increased resolution until BEC subtypes could be differentiated (resolution = 1.2) and then collapsed small clusters (<0.5% of cells) into their nearest neighbor cluster. This allowed us to identify cell supertypes as the highest resolution of clusters that all individuals (except those with exceptionally low cell counts) contributed to (Figure S2; Table S4).

We used Azimuth [37] to annotate cells at the subclass and supertype levels using published reference atlases (using source code from the satijalab/azimuth github) (Figures 1C-I, S2; Table S4). Specifically, we preprocessed the data (*SCTransform* function), found anchors between the query and reference (*FindTransferAnchors* function), gathered all levels of annotation (*TransferData* function), calculated the embeddings of the query data on the reference SPCA (*IntegrateEmbeddings* function), calculated the query neighbors in the reference with respect to the integrated embeddings (*FindNeighbors* function), corrected the Neighbors to account for the downsampling (*NNTransform* function), projected the query to the reference UMAP (*RunUMAP* function), and calculated the mapping scores (*MappingScore* function) [37] (see *myAzimuth* function in github for specific arguments). We used multiple reference atlases, including two from the Allen Institute (M1 – 10X GENOMICS, 2020; MULTIPLE CORTICAL AREAS – SMART-SEQ, 2019) [100,101] and the Human Brain Cell Atlas v1.0 (using cell type-specific datasets in CELLxGENE) [38]. Gene x cell count matrices (and associated metadata) for these atlases were downloaded and then prepared for use in Azimuth by: i) preprocessing the data (removing outliers and running *SCTransform*, *RunPCA*, and *RunUMAP*); ii) making the Seurat object compatible with Azimuth (*AzimuthReference* function); and iii) validating the references (*ValidateAzimuthReference* function). For each reference dataset, annotations for each supertype were visualized as the proportion of cells in each cluster assigned to a given cell type (*prop.table* function).

### Differential cell type proportions

We tested for sex differences in cell type proportions both within and across brain regions (Figure 2A; Table S5). In both cases, we calculated cell type proportions based on cell type and sample information and performed a variance stabilizing transformation (arcsin-square root transformation) on the proportions (*getTransformedProps* function in the R package *speckle* [40]). Within each region, we modeled these normalized proportions as a function of sex, age, and sample-level technical effects (PMI, RIN, batch, total number of reads, total number of cells) (*propeller.ttest* function in the R package *speckle* [40], which calls *lmFit* and associated functions from *limma* [102]). Across regions, we included individual as a random effect and added region as a fixed effect. We first estimated the within individual correlations (*duplicateCorrelation* function in the R package *limma*; specifying individual as the blocking variable) and specified the average estimated within individual correlation in the linear model (using *lmFit* and associated functions from *limma*).

### Pseudobulk aggregation, normalization, and filtering

We created pseudobulks by aggregating counts across all cells (*ADPBulk* function in *adpbulk*) for: i) each sample; and ii) each subclass x sample. Counts were summed across technical replicates of the same biological samples (Table S1). We normalized the read count matrices (for each per sample x subclass pseudobulk) using the functions *calcNormFactors* (R package edgeR) [103] and *voom* (R package *limma*) [102]. Prior to further RNA-seq data analysis, we filtered out genes that were very lowly or not detectably expressed in our samples. Specifically, within each brain region and subclass, we removed any gene with mean CPM<5 across all samples. This procedure resulted in a mean of 11,170 genes (range: 7,880-12,322; fewest in Oligo, most in VIP inhibitory neurons), and 18,263 unique genes were detectably expressed in at least one brain region and subclass. These data (normalized log2 counts per million reads) were used throughout the statistical analyses described below.

### Variance partitioning

We performed variance partitioning on the normalized, filtered pseudobulk expression matrices using the *fitExtractVarPartModel* and *plotVarPart* functions in the R package *variancePartition* [104,105]. This function allowed us to fit a linear mixed model to estimate contribution of multiple sources of variation while simultaneously correcting for all other variables. For analyses of overall expression variance, we used all per sample and supertype pseudo bulk matrices and modeled the expression of each gene as a function of cell type, brain region, individual, sex, age, RIN, PMI, # reads, # cells. For analyses of variance within each subclass and brain region, we modeled the expression of each gene (for all pseudobulk samples of the same cell type and region) as a function of # cells, age, RIN, PMI, sex, # reads. Sex was modeled as a random effect in the overall model and as a continuous factor in the cell type x region models (female = −1, male = 1) [105]. We extracted and visualized the fraction of variance explained by each biological or technical term, in addition to the residual variance (Figure 2B; Table S6). We also visualized the proportion of variance explained by sex for each gene (within each cell subclass and region) (Figure S3; Table S7).

### Differential expression analysis

We initially performed differential expression analysis of pseudobulk samples within each region and subclass. We only included samples with at least 30 cells for a given cell type. Prior to downstream analyses, mitochondrial and ribosomal genes were removed. We then identified outlier counts using Cook’s distances and replaced these values with the trimmed mean over all samples (*replaceOutliers* function in the R package *DESeq2* [106]). We calculated scaling factors using the trimmed mean of M-values method [107] (*calcNormFactors* function in the R package *edgeR* [103]; method = ‘TMM’). We transformed the data to normalized log2 counts per million using *voomWithQualityWeights* (R package *limma* [102]), which combines observational-level weights from *voom* (derived from the mean-variance relationship) with sample-specific weights [108]. We performed linear modeling to estimate sex differences in expression using relevant functions in the R package *limma* (functions *lmFit*, *makeContrasts*, *contrasts.fit*, and *eBayes*). For each gene (in each cell type and region), the normalized expression level was modeled as a function of sex, age, PMI, RIN, total number of cells, and total number of reads. Continuous variables were centered and scaled prior to modeling (*scale* function in the R package *base*).

Following other recent studies [12,45], we then performed a meta-analysis of sex effects across all regions and cell types by applying multivariate adaptive shrinkage (R package *mashr* [43]). This approach uses an empirical Bayesian approach to estimate correlations among different conditions (i.e., unique regions x cell type combinations), using the outputs of the voom-limma models (i.e., the per gene βs and their standard errors) described above as priors. For missing data, βs were set to 0 and standard errors were set to 1e6. We estimated canonical (*cov_canonical* function) and data-driven covariances. The latter were identified using strong signals identified by running a condition-by-condition analysis on all the data (*mash_1by1* function) and extracting those results with local false sign rate (LFSR) < 0.2 in any condition. We performed both PCA (*cov_pca* function; with the number of PCs set to N = 19 cell types) and Empirical Bayes Matrix Factorization (EBMF) (*cov_flash* function; constraining the factors to be nonnegative) and used these to initialize extreme deconvolution (*cov_ed* function; max iterations = 10). We used both canonical and data-driven covariances to fit the mash model and estimated residual correlations using the EM algorithm (*mash_estimate_corr_em* function; max iterations = 5) (Figure S4). We also considered four additional mashr models (PCA-derived data driven co-variances + EM correlations (no EBMF), PCA + simple correlations (*estimate_null_correlation_simple* function), simple or EM correlations without data driven co-variances) and compared the likelihoods of all models (*get_loglik* function) – the results we present are from the best fit model. Sex-biased genes were defined as those with local false sign rates (LFSRs) < 0.05 (Table S9). Results were similar across pre-mashr (i.e., voom-limma) and post-MASH analyses, such that both sex effects (β_mashr_ and β_voom-limma_) and the numbers of sex-biased genes were correlated across region x cell type combinations (across all β: ρ = 0.513, p < 2.2e-16; excluding the Y chromosome and *XIST*: ρ = 0.511, p < 2.2e-16; across all sex-biased genes, either p_adj_ [voom-limma] or LFSR [MASH] < 0.05; ρ = 0.444, p = 2.01e-12) (Figure S4). As expected, MASH increased power, as more sex-biased genes were detected post-MASH (Figure S4) (see also [12]). Of the N=59,131 total significant (LFSR < 0.05) sex-biased gene x cell type x region combinations in MASH, N = 2,224 (3.8%) were also sex-biased in voom-limma (p_adj_ < 0.05 and LFSR < 0.05), N = 50,381 (85.2%) had the same sign but non-significant sex-bias in voom-limma (p_adj_ > 0.05 and LFSR < 0.05), N = 6,526 (11%) had the opposite sign but non-significant sex-bias in voom-limma (p_adj_ > 0.05 and LFSR < 0.05), and 0 (0%) showed significant sex-biases in the opposite directions (Figure S4).

We then performed functional enrichment separately on ubiquitously male-biased (N=58) and female-biased (N=61) (Table S13) using g:Profiler [109] (background = “all known genes”) and identified enriched terms as those with g:SCS values < 0.1 (Table S14). g:SCS method is the default method for computing multiple testing correction for p-values gained from GO and pathway enrichment analysis. It corresponds to an experiment-wide threshold of a=0.05, i.e. at least 95% of matches above threshold are statistically significant.

### Transcriptome wide impact

We also estimated the transcriptome-wide impact of sex (TWI_sex_) for each region and subclass using TRADE [30] (Figures 2D, S6; Table S11). Similar to mashr, this approach uses the outputs of the voom-limma models (i.e., the per gene βs and their standard errors) as inputs. TRADE fits a mixture model to the distribution of effect size estimates (βs), incorporating standard errors to account for sampling variation. TWI is estimated as the variance of the effect size distribution, and is a measure of the overall transcriptomic change. Large TWI values may be obtained when there are large effects on a few genes and/or when there are smaller effects on many genes. The significance of TWI was estimated in two ways: 1) jackknife standard error (which measures our confidence in the magnitude of TWI); and ii) permutation (which measures our confidence that TWI is not zero).

### Subsampling analysis

To address the varying numbers of nuclei across cell subclasses and samples, we also re-estimated the sex effects on gene expression (β_mashr_ and LFSRs) and sexTWI after randomly selecting 50 cells from each sample for each major cell type. This subsampling and testing was repeated 5 times for sex effects on gene expression (β_mashr_ and LFSRs) and 20 times for sexTWI estimates. We then correlated region x cell type rankings – for both the numbers of sbGENEs and sexTWI values – between our original findings and the median rank across all iterations (Figures S6, S7).

### Allele specific expression of X chromosome genes

To identify cases of escape from XCI in XX individuals, we used established approaches for estimating allele specific expression (ASE) in single cell data [54] and using these data to identify escape genes [110].

To call and annotate SNPs, we followed the preprocessing pipeline for scLinaX [54]. We first subset our CellRanger (v5.0.1) [90] bam files of all samples to chrX using *fetch* command. We then genotyped chrX for all single nuclei using *cellsnp-lite* [111] by: i) first running *cellsnp-lite* in Mode 2b to detect SNPs and call heterozygous variants; and then ii) running *cellsnp-lite* in Mode 1a to genotype single cells, using called heterozygous variants from *cellsnp-lite* Mode 2b. We processed the *cellsnp-lite* outputs using a custom R script from the scLinaX User Guide (RCODE_process_cellsnp.r) [54]. We functionally annotated genetic variants outputs using *table_annovar.pl* in *ANNOVAR* [112] (hg38), including gene-based annotations (“refGene” = NCBI RefSeq, “ensGene = Ensembl gene models) and filter-based annotations (“avsnp150” = dbSNP build 150, “ALL.sites.2015_08” = 1000 Genomes Project). Variant annotation was performed using a custom function that reads in *ANNOVAR* output, constructs a unique SNP identifier (chr:position:ref:alt), and extracts relevant annotation fields, including gene region (Func.refGene), gene name (Gene.refGene), and allele frequency in the 1000 Genomes Project (ALL.sites.2015_08). Missing allele frequency values were assumed to be 0. Annotated SNPs were then merged with the RNA SNP dataset by this unique identifier. To ensure specificity, we excluded variants associated with multiple genes and retained only those mapped to informative genic regions (intronic, UTRs, exonic, ncRNA exonic/intronic, and splicing). Finally, to focus on common variants, we filtered out SNPs absent from the 1000 Genomes Project (i.e., ALL_Freq=0).

To assess allelic expression at the single cell level, we used a modified version of the approach by [110]. We first aggregated SNP-level counts per cell by grouping the filtered dataset by cell_barcode, SNP_ID, and Gene, and summing reference and alternate allele counts. The total read count per SNP (tot) was calculated as the sum of reference and alternate counts, and the reference allele ratio (ar) was computed as ar = ref / tot. SNPs were classified as biallelic if the reference ratio was between 0.1 and 0.9; otherwise, they were labeled as monoallelic [110]. Only SNP-cell combinations with at least two total supporting reads (tot>=2) were included. We removed cells with biallelic expression of *XIST*, which would suggest a doublet. To quantify gene-level evidence of biallelic expression, we aggregated SNP data by gene. For each gene in each cell type, we calculated the total number of SNPs (tot_n_snps), the number of biallelic SNPs (bi_n_snps), and the total number of biallelic reads (tot_bi_reads). A binary flag (any.evidence) was assigned to indicate whether a gene had any biallelic SNPs. We also assessed the extent of biallelic expression across cells by counting, for each gene and cell type, the number of unique cells containing the gene (total_cells), the number of cells with at least one biallelic SNP (cells_with_bi), and the number of cells showing exclusively monoallelic expression (cells_only_mono). To ensure reliable estimates, we retained gene x cell type x sample combinations with more than two independent SNPs (tot_n_snps>2). The number of cells with at least one biallelic SNP (cells_with_bi) was then summarized within sexes and cell types (providing the total # of cells with evidence for biallelic expression in each sex and cell type).

To verify that this approach has a low false positive rate for calling biallelic expression at the gene level, we applied it to males and found that 81% of cell type x gene combinations (for nonPAR chrX genes) showed monoallelic expression in males. Biallelic observations in males were primarily due to nine genes with low mappability and high likelihood of mapping errors (due to e.g., pseudogenes, highly similar paralogous sequences, or highly repetitive/low complexity sequences) (*FRMPD4*, *DMD*, *PTCHD1*-*AS*, *IDS*, *TBL1X*, *DANT2*, *FTX*, *IL1RAPL1*, and *LINC00632*) – these were removed from our analysis (increasing the proportion of cell type x gene combinations with monoallelic expression in males to 92%). Having benchmarked our algorithm for calling biallelic expression, we applied it to females. We also retained only gene x cell type combinations where females had more biallelic cells than males for nonPAR chrX genes. We visualized the log-transformed number of biallelic cells (log(N biallelic cells +1)) per cell type and sex for both PAR and nonPAR chrX genes (Figure S9; Table S16).

We compared the escape genes identified in our analyses to those that have been previously reported [50,51] and found N=76 genes that were not reported to exhibit consistent or variable escape in these studies (Figure 2I; Table S16). We then performed functional enrichment on this set of 76 genes using g:Profiler [109] (background = “only annotated genes”) and identified enriched terms as those with g:SCS values < 0.1 (Figure 2I; Table S17).

### Block modelling

We identified clusters of region x cell type combinations with similar sex effects. Specifically, for all N = 114 region x cell type pairwise combinations, we calculated the Spearman rank order correlation using all available pairs of autosomal genes (*cor* function in the R package *WGCNA* [113]) (Table S18). This 114×114 correlation matrix was then clustered using Weighted Stochastic Block Modelling (WSBM) (*BM_gaussian* function in the R package *blockmodels* [114]) to define sets of region x cell type combinations that show coordinated sex effects on expression. WSBM is a generative algorithm that can detect both assortative and non-assortative classes of cluster solution. The optimal number of clusters was derived from the clustering solution that maximized the integrated classification likelihood (ICL) [115]. Clusters were given descriptive names based on their content. Correlation matrices were visualized using a heatmap from the R package pheatmap [116] and a network visualization of the WSBM clustered correlation matrix was created using the R package *igraph* [117] (only correlations >0.3 were visualized) (Figures 3A, S10; Table S19).

### Linking pairwise sex effects to cortical features

We obtained values for ten brain maps across Glasser parcellation areas [118] from Sydnor and colleagues [63]. These maps represented a range of cortical features: allometric scaling (low to high), cerebral blood flow (low to high), cortical thickness (thinner to thicker), evolutionary expansion (low to high), externopyramidisation (externo-to interno-pyramidal), functional hierarchy (unimodal to transmodal), gene expression (sensory to association area expression), Neurosynth categories (motor/perception to thought/emotion), aerobic glycolysis (low to high), and anatomical hierarchy (low to high T1/T2 ratio). We performed principal component analysis (PCA) across all Glasser areas and brain maps (*prcomp* function in R package *stats*), extracting five PCs that each explained ≥5% of the total variance (Figures 3B, S11). We examined the loadings of each PC to identify the contributions from the original brain maps (Figures 3B, S11). To relate these PCs to our data, we mapped the six regions from our study onto the Glasser parcellation [118]: AngGyr (TPOJ3), CauIns (52), FusGyr (FFC), IntParS (AIP), InfLTO (LO2), and RetCor (RSC) (Figure S1C). For each PC, we calculated pairwise similarity between all region pairs by: i) computing the absolute difference in PC values between each region pair; and ii) subtracting these values from the maximum observed difference for that PC, yielding a measure of relative similarity (larger values = more similar) (Figures 3B, S11). Finally, for each PC and each cell type, we computed the Spearman rank correlation (*cor* function in the R package *stats*) between the PC-based region similarity values and the pairwise sex effect similarity measures (described in the ‘Block modelling’ section above) (Figures 3B, S11).

### Functional enrichments for sex-biased autosomal gene clusters

We performed gene ontology enrichment analyses on union sets of autosomal genes that were exclusively male-biased or strictly female-biased across all region x cell type combinations in each cluster (see ‘Block modelling’ section) using the *enricher* function in the R package *clusterProfiler* [119]. In each case, the background was the union set of autosomal genes expressed among the region x cell type combinations in a given cluster, and we tested for enrichment of biological processes (BP) and cellular compartments (CC). Significantly enriched terms were those with nominal p < 0.05 (Tables S20-S21). These significant terms were subsequently ranked according to their fold enrichments (i.e., the ratio of the frequency of input genes annotated in a term to the frequency of all genes annotated to that term) within each cluster and the top three terms within each cluster were visualized (Figure 3C).

### Transcription factor enrichments for sex-biased autosomal gene clusters

We used the *findMotifs.pl* function in HOMER (Hypergeometric Optimization of Motif EnRichment) [67] to analyze the promoters of genes that were exclusively male-biased, exclusively female-biased, or sex-biased (male- and/or female-biased) in each cluster (see ‘Block modelling’ section). We searched for motifs from −1000 to +300 relative to the transcriptional start site (TSS). The program assigns weights to the background promoters based on the distribution of GC content in the target gene promoters to ensure that comparable numbers of low and high-GC promoters are analyzed. It also performs auto-normalization to remove sequence content bias from lower order oligos (1/2/3-mers) by adjusting background weights based on the target distribution. The hypergeometric distribution is used to score motifs. We used both HOMER’s curated set of >400 known vertebrate motifs (-mset vertebrates) and the HOCOMOCO v11 human full collection (primary and alternative binding models, p<0.001) [66]. We extracted results for the sex chromosome and steroid hormone-related motifs from each set (listed below). Sex chromosome motifs were filtered to genes expressed in any condition in our dataset. P-values were adjusted within each cluster, TF set, and sex-biased gene set (male-biased or female-biased) across TFs using the Benjamini-Hochenberg method [120] (*p.adjust* function in the R package *stats*). Motifs with p_adj_<0.1 were considered significantly enriched in the promoter regions of the target gene set (Figures 3D; Table S22).

Homer TF set:

- Hormone-related

- ARE(NR)/LNCAP-AR-ChIP-Seq(GSE27824)/Homer = androgen receptor
- ERb(NR),IR3/Ovary-ERb-ChIP-Seq(GSE203391)/Homer = estrogen β receptor
- ERE(NR),IR3/MCF7-ERa-ChIP-Seq(Unpublished)/Homer = estrogen α receptor
- PGR(NR)/EndoStromal-PGR-ChIP-Seq(GSE69539)/Homer = progesterone receptor
- PR(NR)/T47D-PR-ChIP-Seq(GSE31130)/Homer = progesterone receptor
- X chromosome

- ZFX(Zf)/mES-Zfx-ChIP-Seq(GSE11431)/Homer
- TFE3(bHLH)/MEF-TFE3-ChIP-Seq(GSE75757)/Homer
- ZBTB33(Zf)/GM12878-ZBTB33-ChIP-Seq(GSE32465)/Homer
- ZNF41(Zf)/HEK293-ZNF41.GFP-ChIP-Seq(GSE58341)/Homer
- ZNF711(Zf)/SHSY5Y-ZNF711-ChIP-Seq(GSE20673)/Homer
- Elf4(ETS)/BMDM-Elf4-ChIP-Seq(GSE88699)/Homer
- Elk1(ETS)/Hela-Elk1-ChIP-Seq(GSE31477)/Homer
- Zic3(Zf)/mES-Zic3-ChIP-Seq(GSE37889)/Homer HOCOMOCO TF set:
- Hormone-related

- ANDR_HUMAN.H11MO.0.A = androgen receptor
- ESR1_HUMAN.H11MO.0.A = estrogen α receptor
- ESR2_HUMAN.H11MO.0.A = estrogen β receptor
- PRGR_HUMAN.H11MO.0.A = progesterone receptor
- X chromosome

- ZFX_HUMAN.H11MO.0.A
- ARX_HUMAN.H11MO.0.D
- ELK1_HUMAN.H11MO.0.B
- FOXO4_HUMAN.H11MO.0.C
- KLF8_HUMAN.H11MO.0.C
- MECP2_HUMAN.H11MO.0.C
- NR0B1_HUMAN.H11MO.0.D
- TAF1_HUMAN.H11MO.0.A
- TFE3_HUMAN.H11MO.0.B
- ZIC3_HUMAN.H11MO.0.B
- ZNF41_HUMAN.H11MO.0.C
- PAR

- ZBED1_HUMAN.H11MO.0.D

### Disease enrichments for sex-biased autosomal gene clusters

We obtained sex-stratified GWAS results from the UK Biobank (round 2) [70] for the following sex-biased phenotypes:

- AD (father) = Illnesses of father: Alzheimer’s disease/dementia (phenotype code: 20107_10)
- AD (mother) = Illnesses of mother: Alzheimer’s disease/dementia (20110_10)
- PD (father) = Illnesses of father: Parkinson’s disease (20107_11)
- PD (mother) = Illnesses of mother: Parkinson’s disease (20110_11)
- OCD = Mental health problems ever diagnosed by a professional: Obsessive compulsive disorder (OCD) (20544_7)
- Schizophrenia = Schizophrenia, schizotypal and delusional disorders (F5_SCHIZO)
- Mania = Mental health problems ever diagnosed by a professional: Mania, hypomania, bipolar or manic-depression (20544_10)
- MS = Diagnoses - main ICD10: G35 Multiple sclerosis (G35)

Then, for each significant SNP in the sex-stratified GWAS datasets (p<5e-05), we identified the nearest gene using the *closest* function in *bedtools* [121]. We tested for overlap between the resulting gene sets and the union of strictly male-biased or strictly female-biased autosomal genes identified in each cluster (see ‘Block modelling’ section) using Fisher’s exact tests (*fisher.test* function in R package *stats*). In each case, the background was the union set of autosomal genes expressed among the region x cell type combinations in a given cluster. Because male- and female-specific GWAS results are often highly similar [88], we estimated the log_2_ ratio of the odds ratios (OR_male-GWAS_/OR_female-GWAS_) for each disease and sex-biased gene set combination. This provided a measure of the relative enrichment for male-vs. female-specific GWAS (ratios > 0 = OR_male-GWAS_ > OR_female-GWAS_; ratios < 0 = OR_male-GWAS_ < OR_female-GWAS_) (Table S23). We visualized these ratios when either OR_male-GWAS_ or OR_female-GWAS_ was significant (nominal p<0.05) (Figures 3E; Table S23). For example, female-biased genes in cluster 4 were more strongly enriched for the female-vs. the male-specific multiple sclerosis (MS) GWAS (OR_female-GWAS_=1.615, p=0.005; OR_male-GWAS_=1.117, p=0.283; OR ratio = log_2_(1.117/1.615) = −0.532; OR_male-GWAS_ < OR_female-GWAS_ so OR ratio < 0) (Table S23), and this OR ratio is shown in Figure 3E.

#### Supplementary Figures

**Figure S1** | Determination of cortical sample locations based on sex differences in cortical volume. (A) Cortical surface projections of significant sex differences (t statistics of males vs. females, family wise error < 0.05) in regional gray matter volume (rGMV) beyond the total GMV that were identified in two independent large-scale datasets from the Human Connectome Project (HCP [122]) and the UK Biobank (UKB [123]) [29]. (B) Six cortical regions (2 male-biased: FusGyr and InfLTO; 2 female-biased: IntParSul and CauIns; 2 unbiased: AngGyr and RetCor) selected on the conjunction of sex differences in rGMV of these two datasets — average t statistics at voxels showing significant sex differences with the same direction in both HCP and UKB. (C) These six selected cortical regions mapped to unique parcels of the standard HCP cortical parcellation [118] — AngGyr (TPOJ3), CauIns (52), FusGyr (FFC), IntParS (AIP), InfLTO (LO2), and RetCor (RSC) on the right hemisphere with different views for demonstration. (D) Images showing dissection of these six selected cortical regions from brain slabs for following nuclei isolation and 10x snRNAseq. Arrow/text colors represent the six selected brain regions, matching Fig. 1A, (unbiased AngGyr = gray; CauIns = blue, FusGyr = green; InfLTO = black; IntParSul = orange; RetCor = purple).

**Figure S2** | UMAP plots for each cell class (excitatory neurons, inhibitory neurons, glial cells). Colors represent cell subclass, cell supertype, brain region, sex, mitochondrial content (“Mito”), UMI counts (“UMI”), and marker gene expression (see titles of individual plots and legends).

**Figure S3** | Boxplots showing the proportion of variance explained by sex within each region x cell subclass combination. Each point represents 1 gene (x’s = X chromosome; triangles = Y chromosome; squares = PAR; circles = autosomal). Colors represent cell subclasses and match those in Figure 1.

**Figure S4** | MASHR results and comparisons to voom-limma results. (A) Counts of male- and female-biased genes in voom-limma (y-axis) versus MASHR (x-axis). Shapes represent direction of sex bias and colors represent cell subclasses (see legend). (B) Figure S4A without ILTO OPC (which shows high numbers of sex-biased genes). (C) Correlations (including R and p-values) between estimated sex effects (β) from voom-limma (y-axis) versus MASHR (x-axis) across all genes in each brain region (columns) and cell subclass (row). (D) Figure S4C without Y chromosome genes and *XIST*. (E) Bar plot showing the estimated mixture proportions for each covariance matrix from MASHR. Canonical covariances (i.e., condition specific effects named for each region x cell subclass combination, equal effects, null, identity, simple_het = correlated effects; see [43] and Methods) and data driven covariances (PCA = covariances produced by singular value decomposition of an overall matrix, “ED_tPCA”; FLASH = covariances produced using Empirical Bayes Matrix Factorization; ED = extreme deconvolution applied; see Methods) are listed along the x-axis. (F) ED_tPCA covariance matrix. (G) ED_PC7 covariance matrix.

**Figure S5** | Distributions of sex effects. (A) Density plots of absolute sex effects in each region x cell subclass combination for autosomal genes. (B) Violin plots of signed sex effects (positive=male-biased; negative=female-biased) in each region x cell subclass combination for autosomal genes. (C) Density plots of absolute sex effects in each region x cell subclass combination for X chromosome genes. (D) Violin plots of signed sex effects (positive=male-biased; negative=female-biased) in each region x cell subclass combination for X chromosome genes.

**Figure S6** | Detailed results for sexTWI. (A) Bar plots of sexTWI values estimated for each region x cell subclass combination. Bar colors represent cell subclasses, box colors represent brain regions (see Figure 1). Grey lines represent jacknife standard errors from leave-one-out estimates. Asterisks (*) represent p-values from permutation tests (null hypothesis: sewTWI = 0). (B) Boxplots of sexTWI within regions and cell classes. (C) Original sewTWI rank versus median rank across 20 subsampling iterations (Methods). Shapes represent brain region, colors represent cell subclasses (see legend).

**Figure S7** | Comparisons between regional sex-biased GMV and measures of sex-biased transcriptomes. (A) Top: Spearman’s ρ values between region-specific sex-biased GMV measures and sexTWI values. sexTWI values were estimated for each region in each cell subclass (rows). “GMV sex bias” = directional GMV sex bias (higher = male-biased, lower = female-biased). Absolute GMV sex bias = absolute value of the GMV sex bias (higher = more sex-biased, lower = less sex-biased). * = nominal p<0.05 (none are significant after adjustment for multiple comparisons). Bottom: Example showing the relationship between region-specific GMV sex bias and sexTWI in L6b excitatory neurons. Dot colors represent brain region (see legend). (B) Top: Similar to Figure S7A (top panel), but for the proportion of expressed genes that are significantly sex-biased (LFSR<0.05; proportion of sbGENEs). Bottom: Example showing the relationship between region-specific GMV sex bias and the proportion of sbGENEs in SST inhibitory neurons. Dot colors represent brain region (see legend). (C) Top: Similar to Figure S7A (top panel), but for pairwise sex effect similarity (i.e., Spearman’s ρ between each pair of brain regions for autosomal gene sex effects in each cell subclass) and pairwise sex-biased GMV measures (i.e., the absolute difference between each pair of regions for each sex-biased GMV measure). Bottom: Example showing the relationship between pairwise sex effect similarity and pairwise GMV sex bias similarity for ADARB2 inhibitory neurons. Dot colors represent each pair of brain regions (see legend).

**Figure S8** | Comparisons of sex effects. (A) sbGENE proportions (relative to the number of expressed genes in a given subclass x region combination). (B) Original sbGENE ranks versus median ranks across 5 subsampling iterations (Methods). Shapes represent brain region, colors represent cell subclasses (see legend). (C) Scatterplots showing sbGENE proportion ranks, versus sbGENE count ranks versus sexTWI ranks. r and p-values are provided. (D) Density plots of gene expression coefficient of variation (CV) within region x cell subclass combinations. (E) Density plots of standard errors (se) for voom-limma sex effects (βvoom-limma) within region x cell subclass combinations.

**Figure S9** | Log(counts+1) of cells with evidence for biallelic expression in males (bottom) and females (top) for nonPAR X genes (left) and PAR genes (right).

**Figure S10** | Network of 13 autosomal clusters (see Figure 3A). Line thickness represents similarity (only correlations >0.3 were visualized).

**Figure S11** | Comparisons of region pairwise similarity for brain maps versus sex effects. (A) approach summary. (B) Correlations between 10 brain maps across all regions. These maps represented a range of cortical features: allometric scaling (low to high), cerebral blood flow (low to high), cortical thickness (thinner to thicker), evolutionary expansion (low to high), externopyramidisation (externo-to interno-pyramidal), functional hierarchy (unimodal to transmodal), gene expression (sensory to association area expression), Neurosynth categories (motor/perception to thought/emotion), aerobic glycolysis (low to high), and anatomical hierarchy (low to high T1/T2ratio). (C) Elbow plot of showing the proportion of variance explained by the top 10 PCs (red dashed line = 5% variance explained). (D) Top left: maps of the top 5 PCs. Top middle: PC loadings for each of the 10 original brain maps across the top 5 PCs. Top right: correlations between PC similarity measures and sex effect similarity measures for the top 5 PCs in each of the 19 cell subclasses. Bottom: scatterplots of the relationships between PC similarity measures and sex effect similarity measures for significant cell subclass x PC combinations (shapes = pairwise region combinations, see legend).

#### Supplementary Tables

**Table S1** | Sample metadata for N=192 samples (all original samples).

**Table S2** | Cell Ranger QC statistics for N=179 samples (low quality samples removed, replicates included separately, see Table S1).

**Table S3** | Alevin QC statistics for N=179 samples (bad samples removed, replicates separate)

**Table S4** | Original cell subclass counts for N=169 samples (low quality samples removed, replicates combined, see Table S1).

**Table S5** | Results from differential proportion analyses (both within and across region analyses are included, see Methods).

**Table S6** | Variance partitioning results showing the proportion of variance attributed to each predictor for each of N = 4,313 genes.

**Table S7** | Variance partitioning regions within regions and cell subclasses (excitatory neurons = columns A-J; inhibitory neurons = columns L-U; other = columns W-AG).

**Table S8** | Variance partitioning correlations (overall = columns A-J, see Table S6; within regions and subclasses = columns M-Q, see table S7).

**Table S9** | MASH-derived sex effects (β_MASH_) for mean gene expression for excitatory neurons (S9A), inhibitory neurons (S9B), and other cell subclasses (S9C). Beta = estimated sex effect; LFSR = local false sign rate.

**Table S10** | Tukey’s HSD results comparing absolute sex effects across the transcriptome (|β_mashr_| regardless of LFSR) between brain regions and cell subclasses. CI = confidence interval.

**Table S11** | sexTWI values for each region x cell subclass combination (permutation p-values = nominal p-values from permutation; jacknife p-values = nominal p-values using jackknife standard errors from leave-on-out analyses).

**Table S12** | Comparisons between regional sex-biased GMV and measures of sex-biased transcriptomes. Spearman’s ρ values and associated p-values are shown. “GMV sex bias” = directional GMV sex bias (higher = male-biased, lower = female-biased). Absolute GMV sex bias = absolute value of the GMV sex bias (higher = more sex-biased, lower = less sex-biased). Within each cell subclass, the regional sex-biased GMV measure was correlated with sexTWI (Columns C-I) or the proportion of expressed genes that were significantly sex-biased (LFSR<0.05; proportion of sbGENEs) (Columns K-Q). Within each cell subclass, we also show correlations between pairwise sex effect similarity (i.e., Spearman’s ρ between each pair of brain regions for autosomal gene sex effects in each cell subclass) and pairwise sex-biased GMV measures (i.e., the absolute difference between each pair of regions for each sex-biased GMV measure) (Columns S-Y).

**Table S13** | Table S9 subset to the N=133 ubiquitously sex-biased genes (i.e., genes that were significantly sex-biased in every region x subclass condition).

**Table S14** | g:Profiler results for ubiquitously male-or female-biased autosomal genes (gene set = column A). MF = molecular function; BP = biological process; CC = cellular compartment; HPA = human protein atlas; HP = human phenotype.

**Table S15** | Tukey’s HSD results comparing the proportions of sex-biased genes (LFSR<0.05) between brain regions and cell subclasses. CI = confidence interval.

**Table S16** | Allele specific expression results for each gene and cell subclass. Columns S-W = summary results for N=136 nonPAR X genes showing evidence for biallelic expression.

**Table S17** | g:Profiler results for N=76 novel XCIe genes (see Table S15). MF = molecular function; BP = biological process; CC = cellular compartment; HPA = human protein atlas; HP = human phenotype.

**Table S18** | Correlations (Spearman’s ρ) of sex effects for autosomal genes (|β_mashr_| regardless of LFSR) across all region x subclass combinations.

**Table S19** | Cluster memberships (columns A-B) and counts of sex-biased (LFSR<0.05) and background genes for each of 13 autosomal gene clusters (columns D-F).

**Table S20** | Biological process (BP) enrichments for male-biased or female-biased genes in each of 13 autosomal gene clusters.

**Table S21** | Cellular compartment (CC) enrichments for male-biased or female-biased genes in each of 13 autosomal gene clusters.

**Table S22** | Motif enrichment results for male-biased or female-biased genes in each of 13 autosomal gene clusters.

**Table S23** | Sex-specific GWAS enrichment results for male-biased or female-biased genes in each of 13 autosomal gene clusters.

## Funding

This work was supported by the Intramural Research Programs of the National Institute on Aging and National Institute of Mental Health.

## Acknowledgements

We thank the Human Brain Collection Core (HBCC) at the National Institute of Mental Health (NIMH) for providing and processing samples.

## Notes

### Competing Interest Statement

The authors have declared no competing interest.

