## Supplementary figures and images for "Sex effects on gene expression across the human cerebral cortex at single cell resolution"

### Fig S1

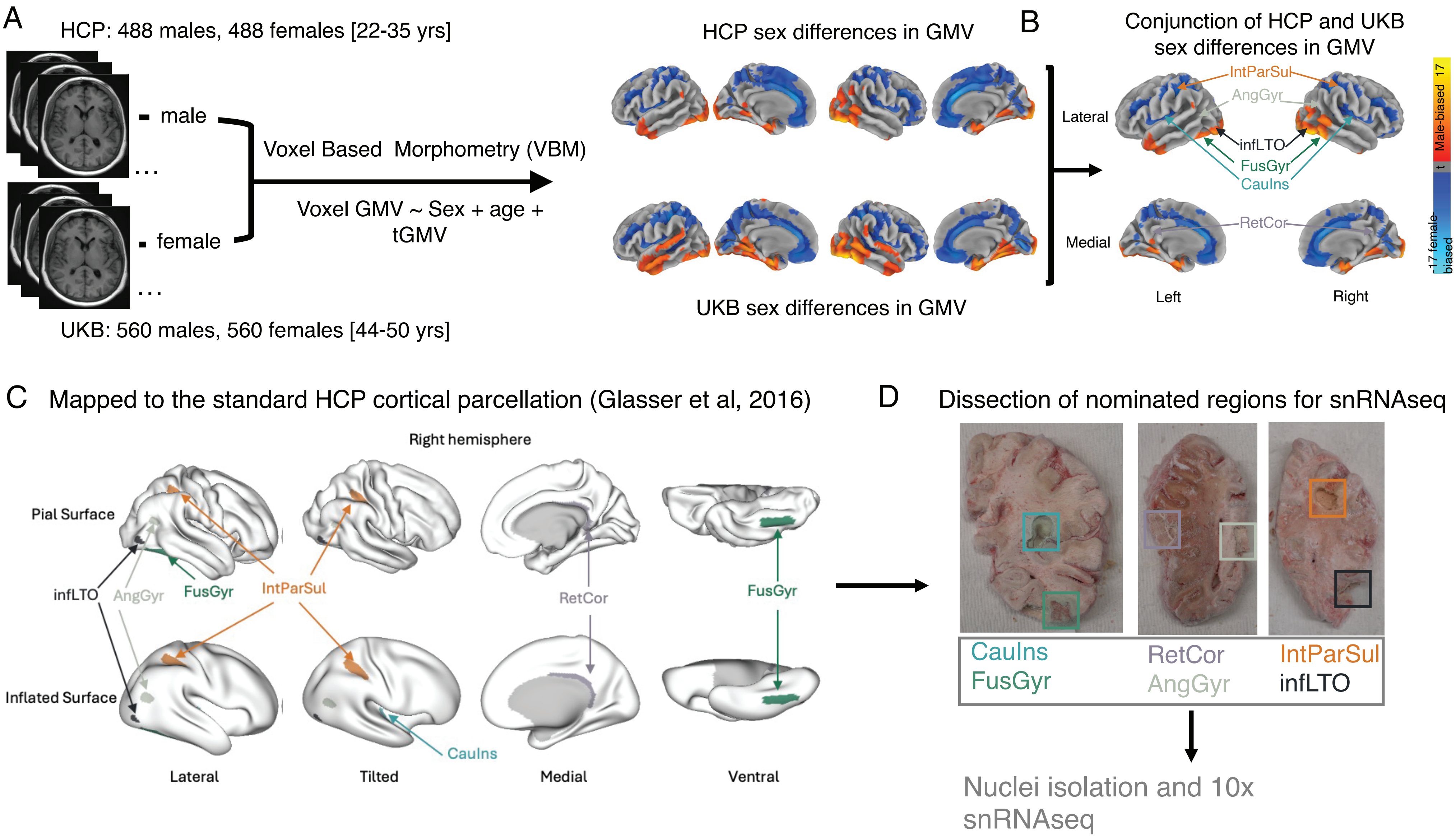

### Fig S2

#### Excitatory neurons

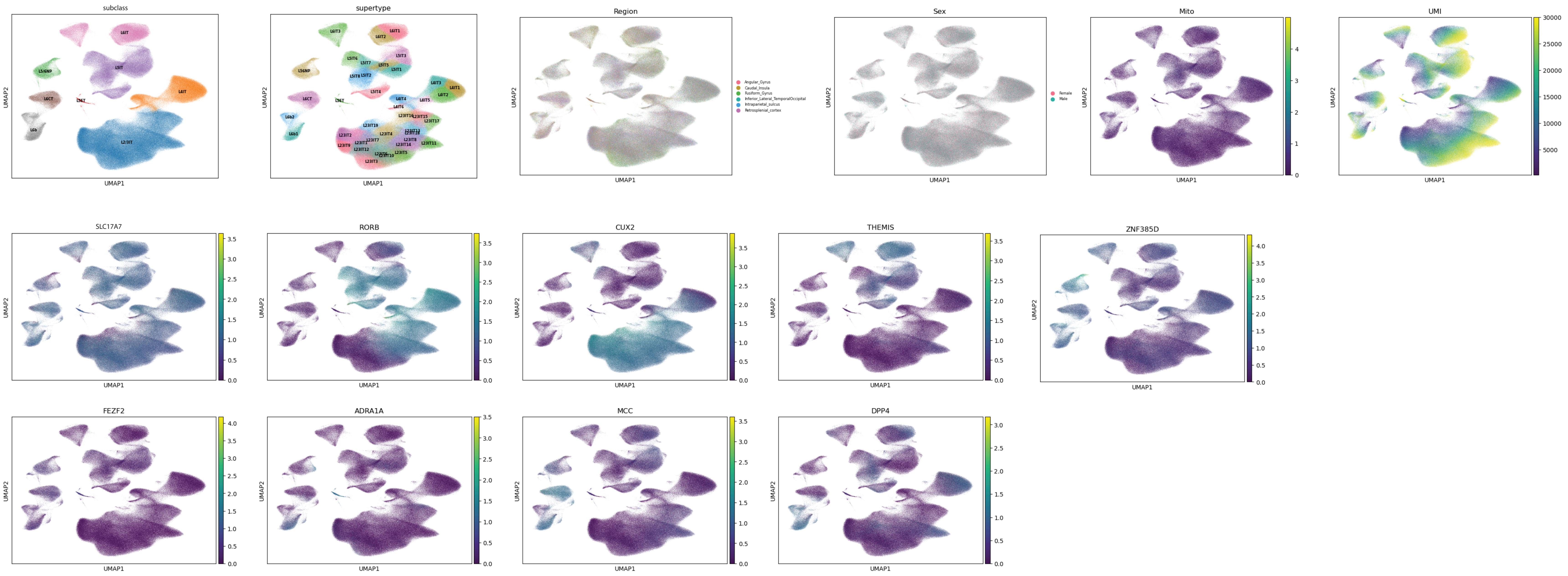


#### Inhibtory neurons

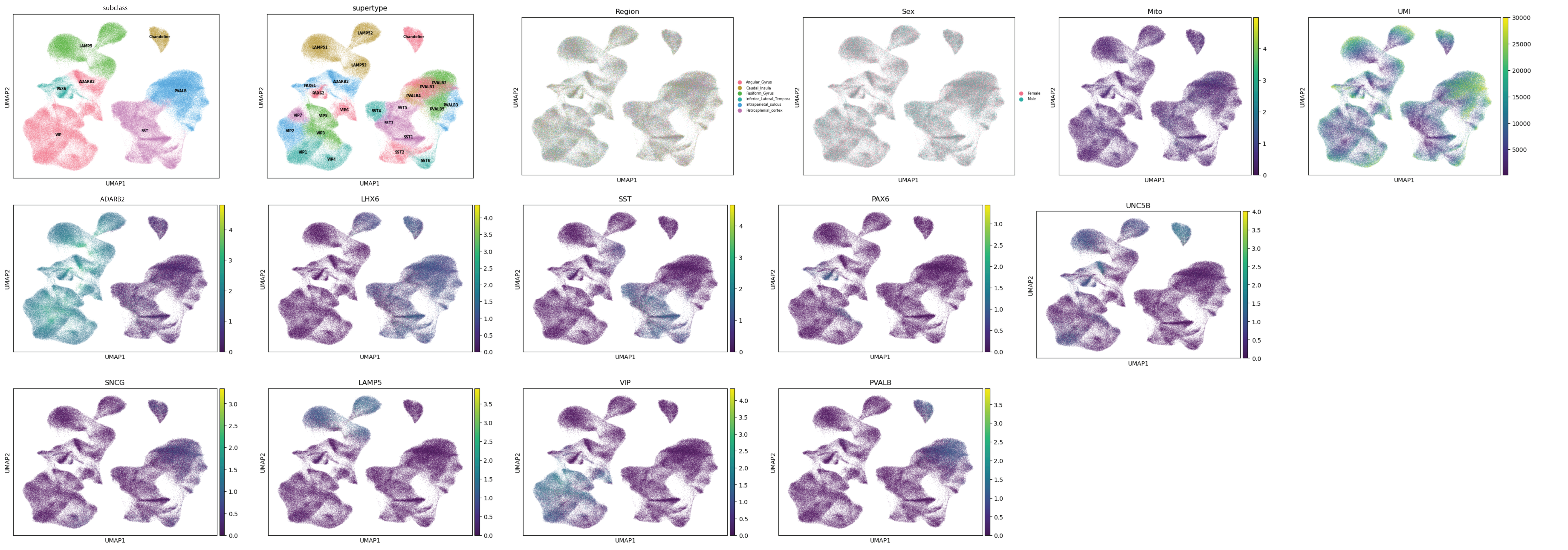


#### Glia

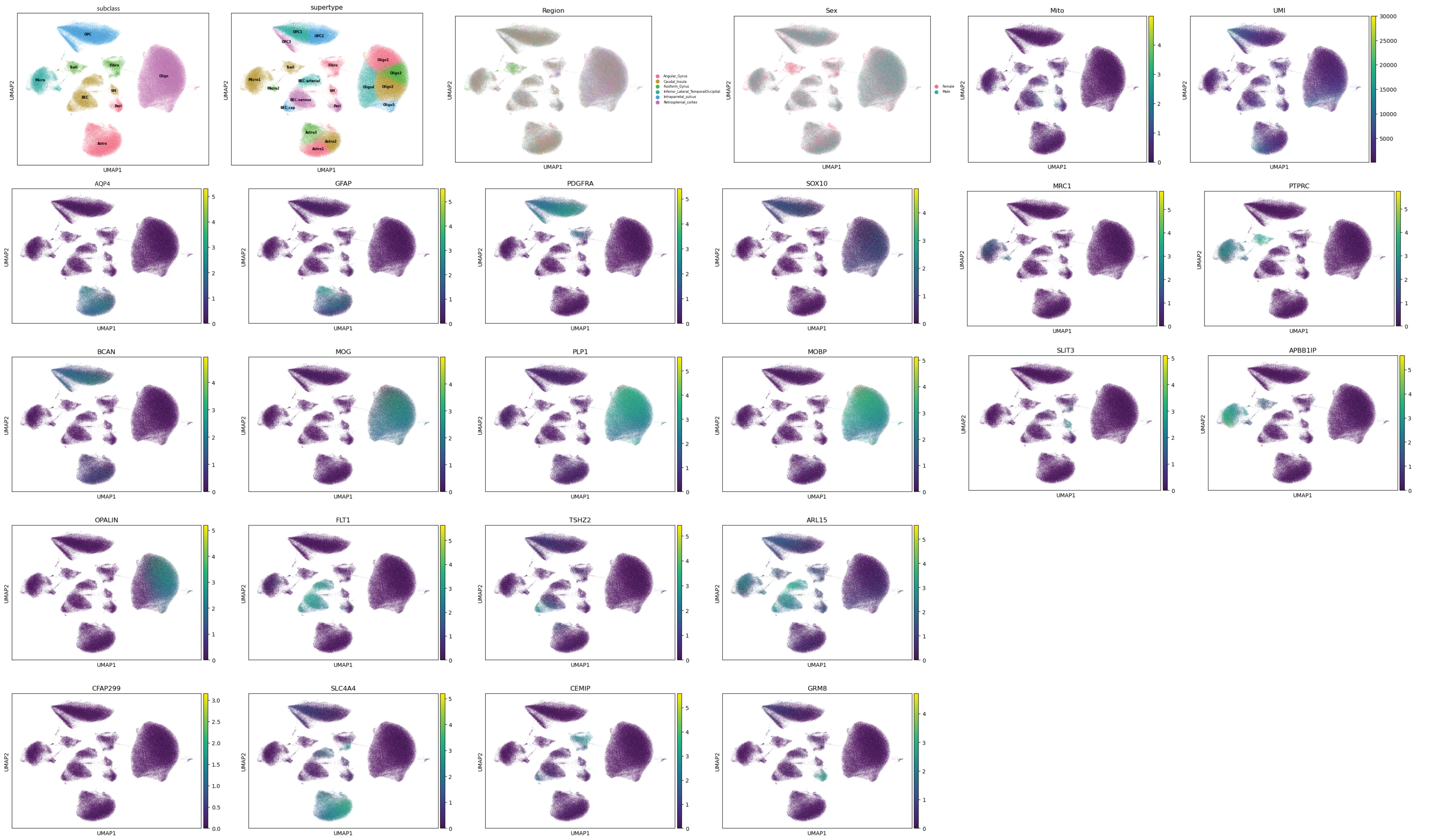

### Fig S3

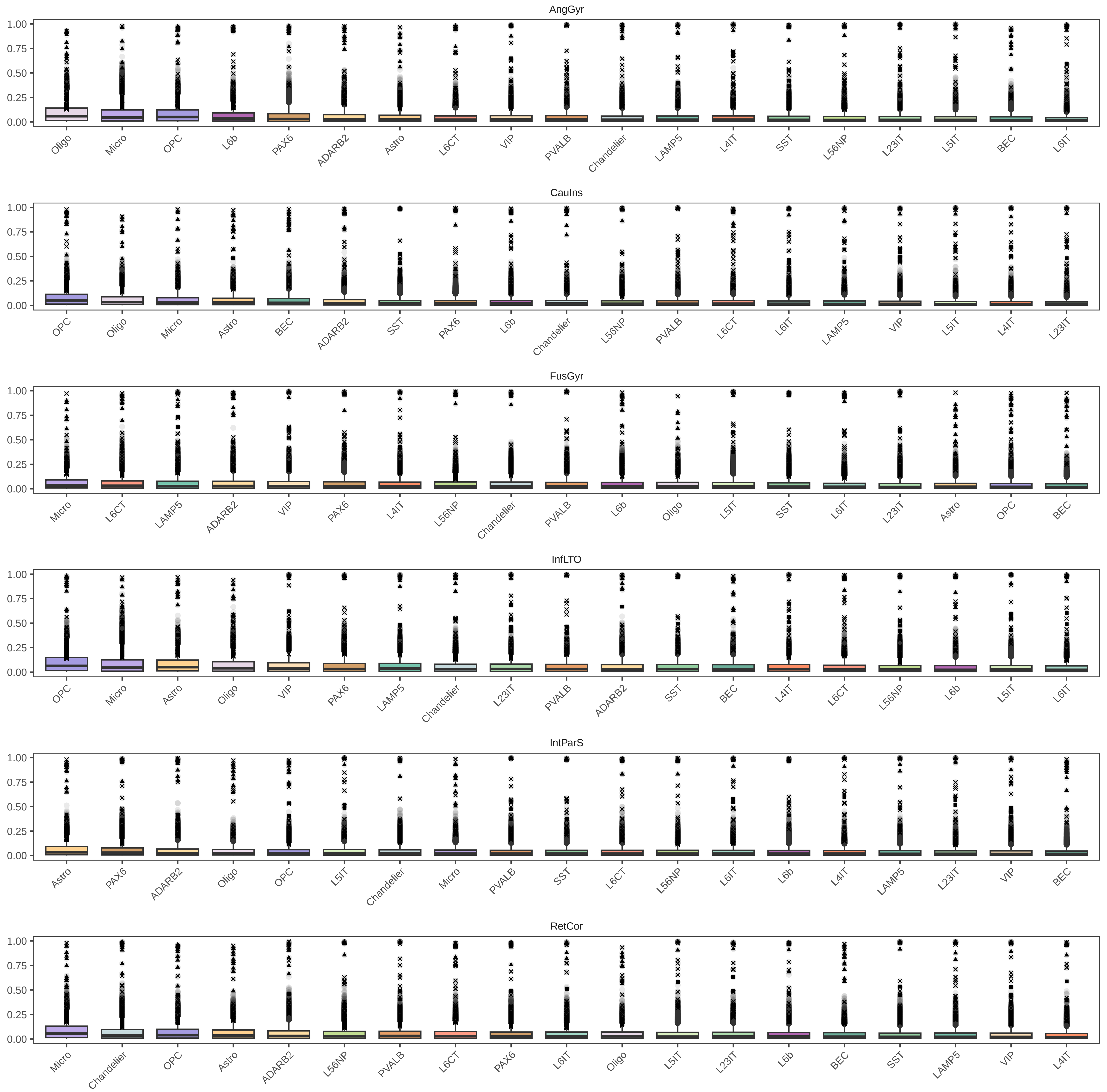

### Fig S4

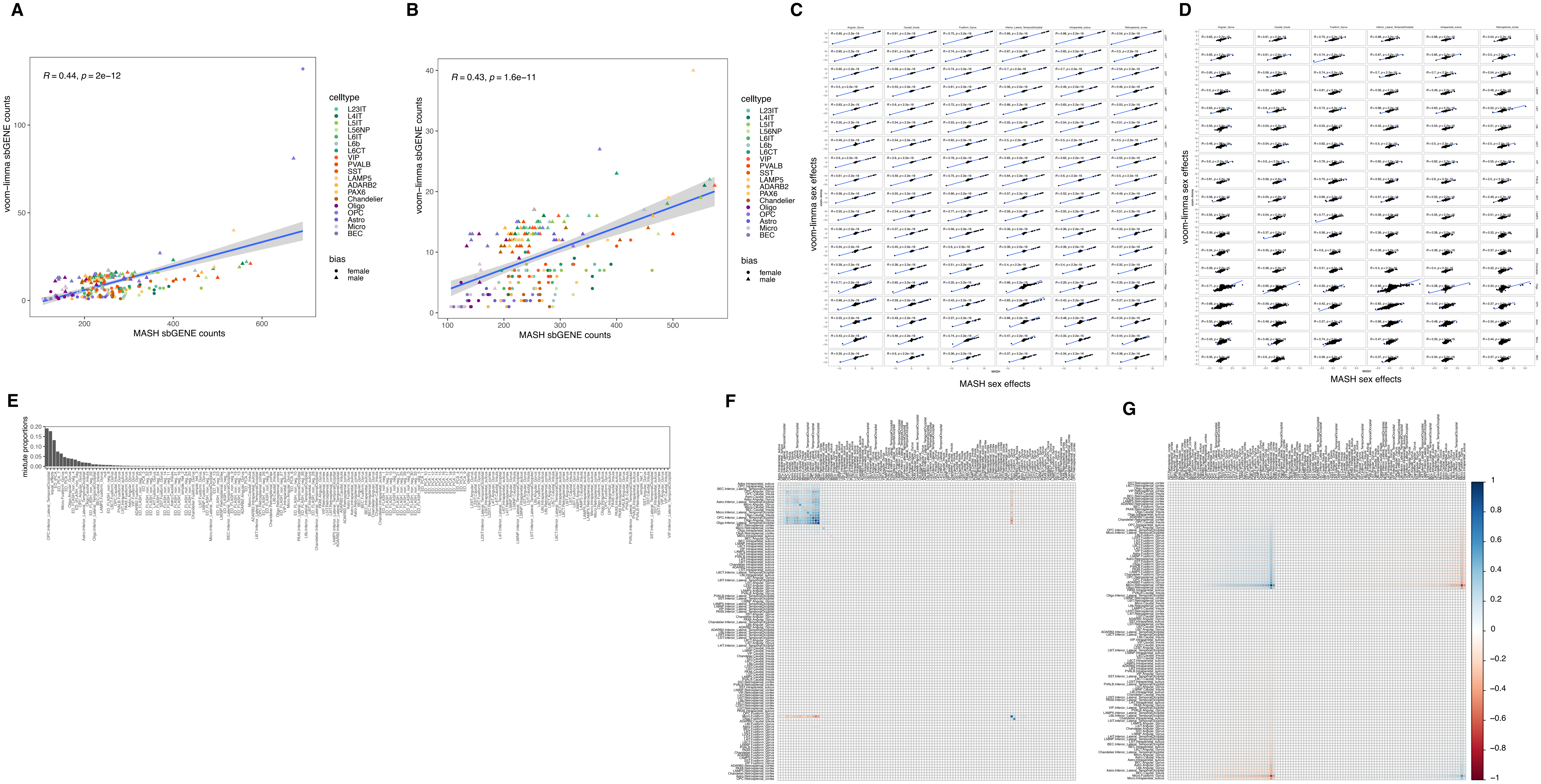

### Fig S5

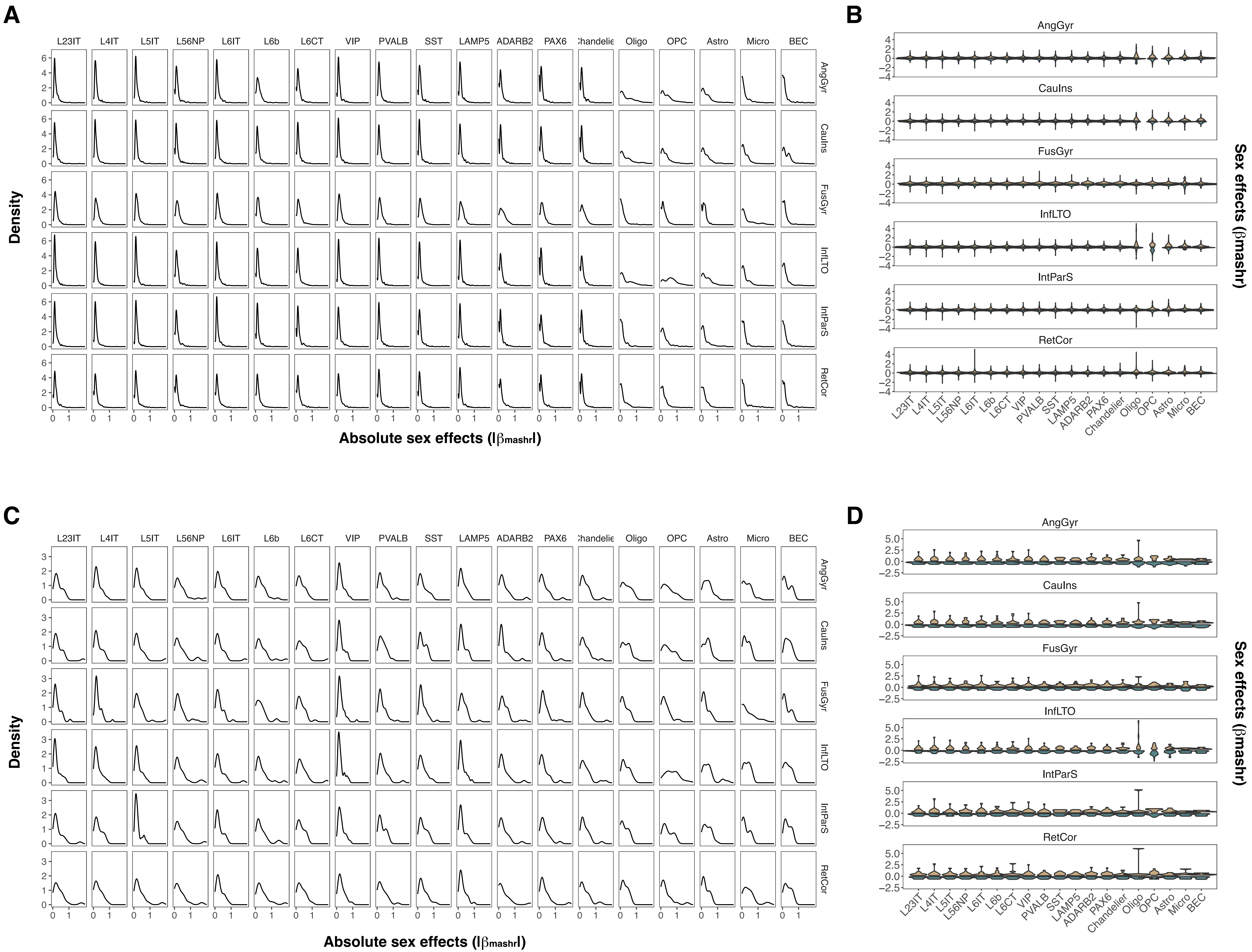

### Fig S6

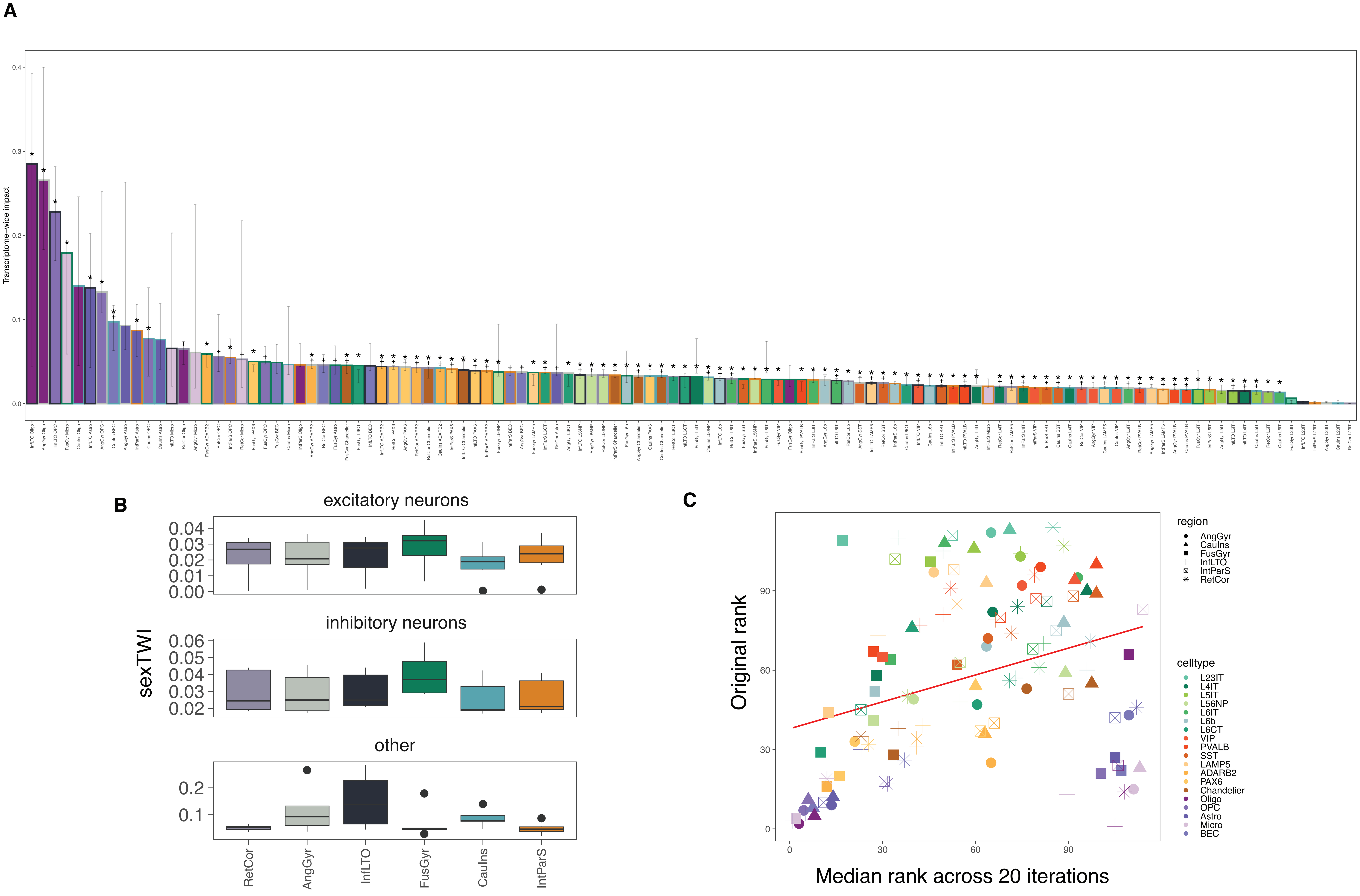

### Fig S7

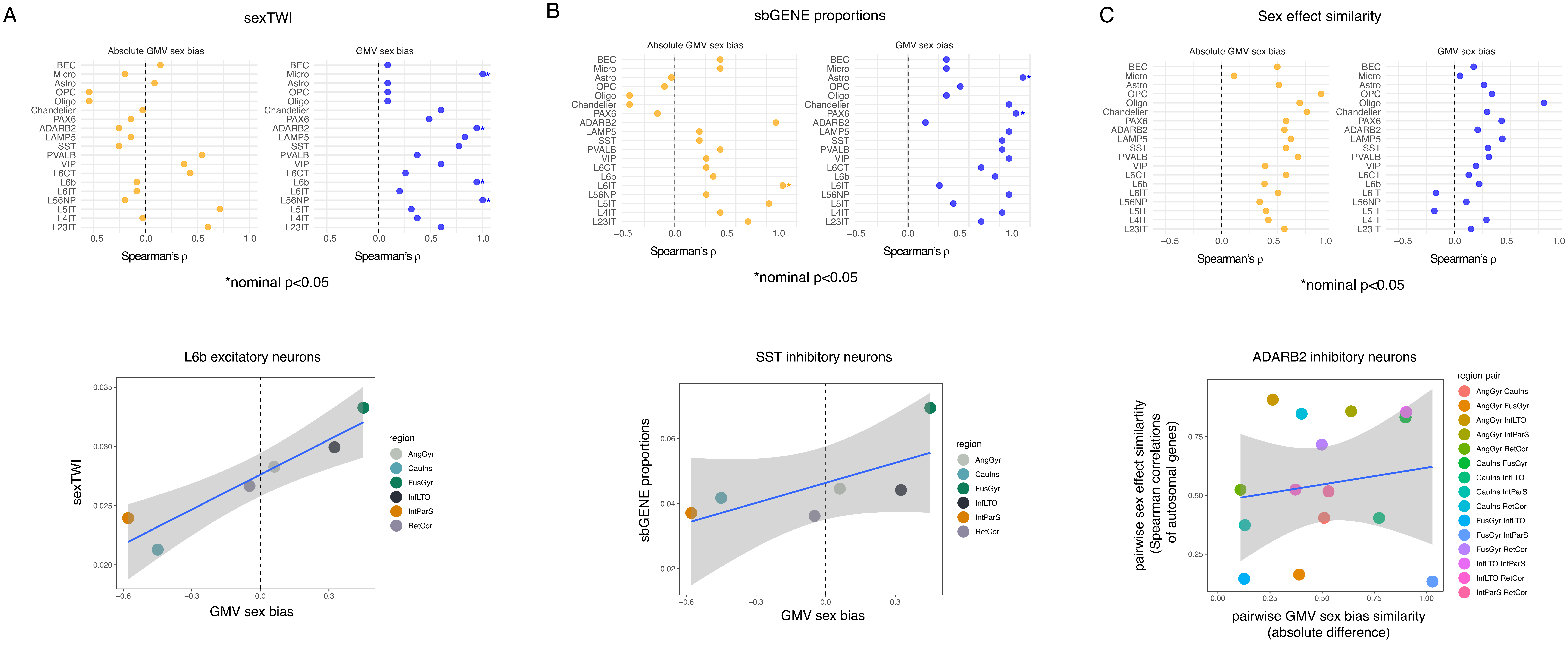

### Fig S8

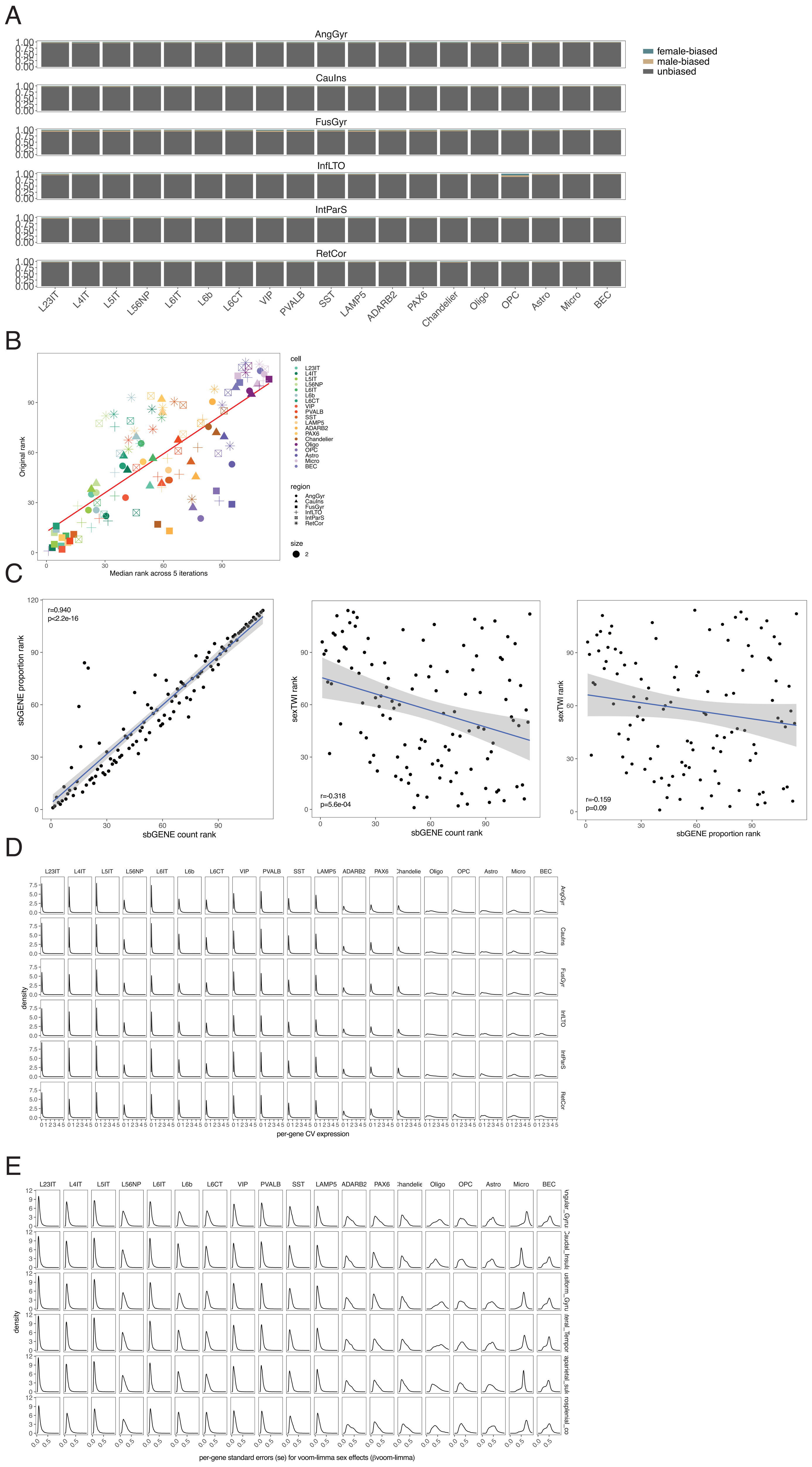

### Fig S9

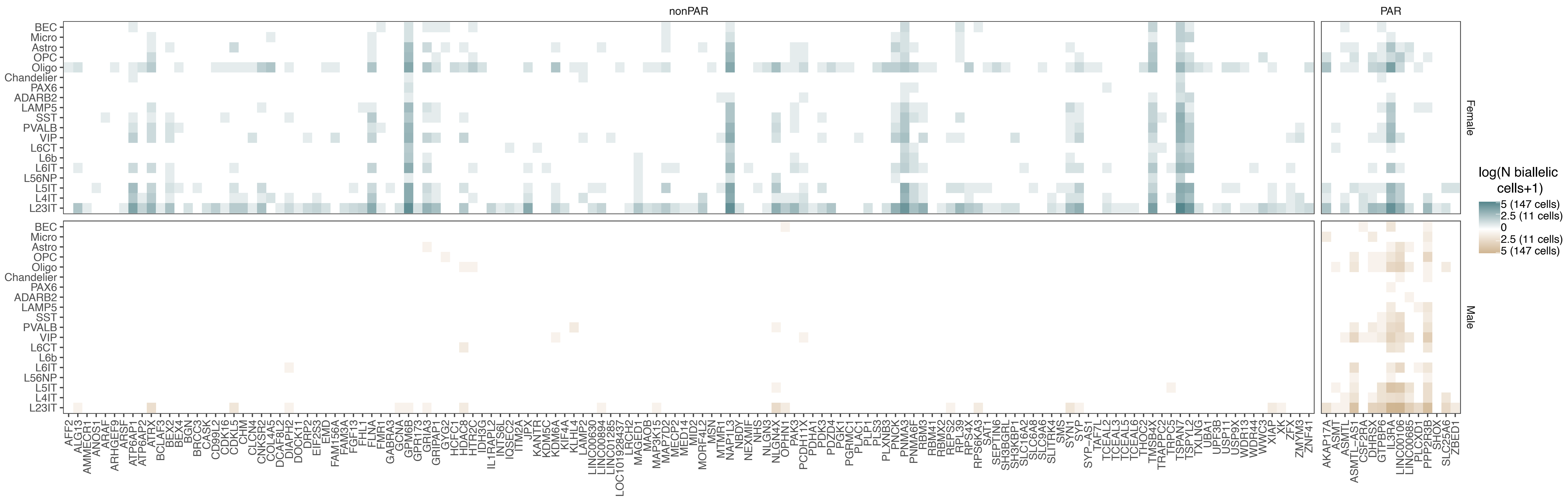

### Fig S10

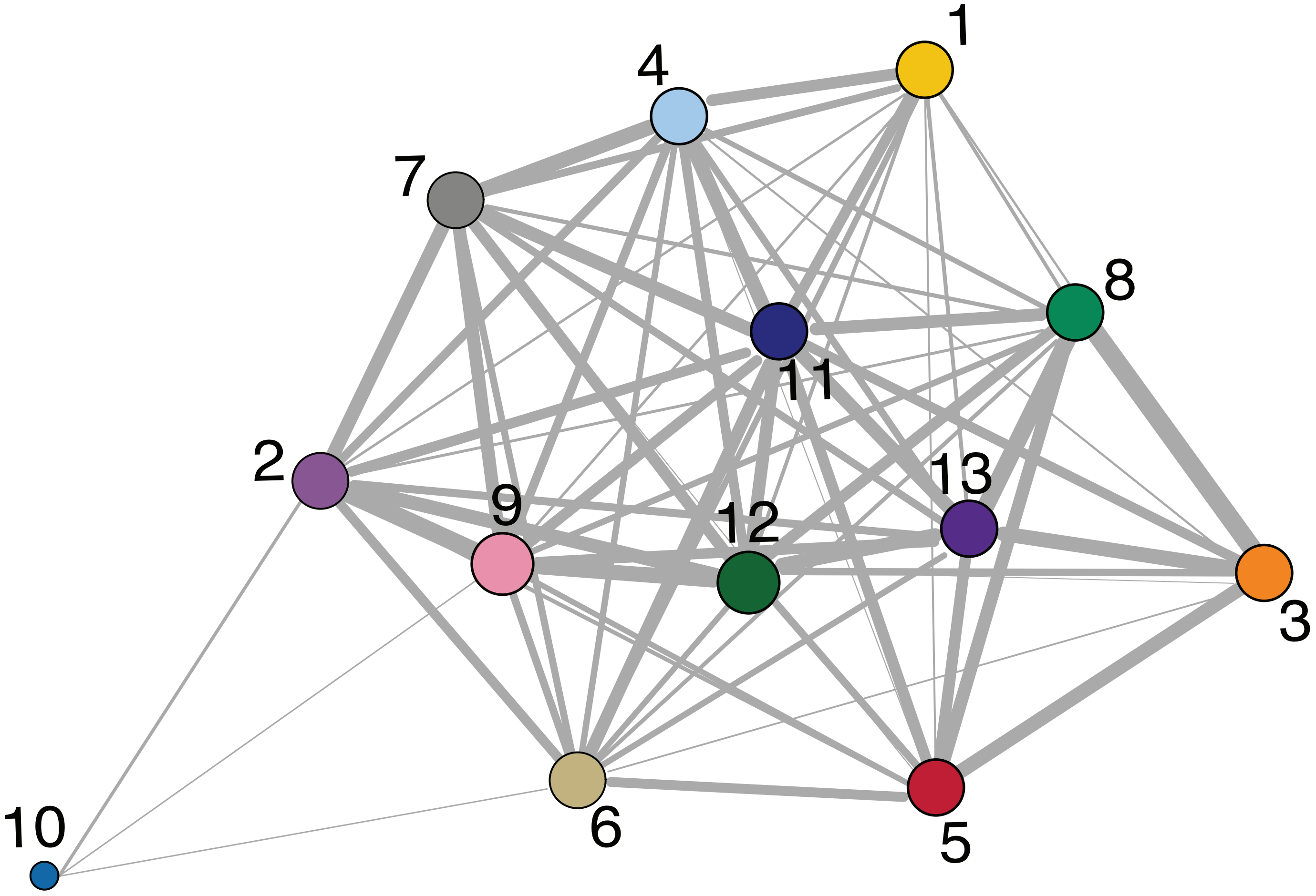

### Fig S11

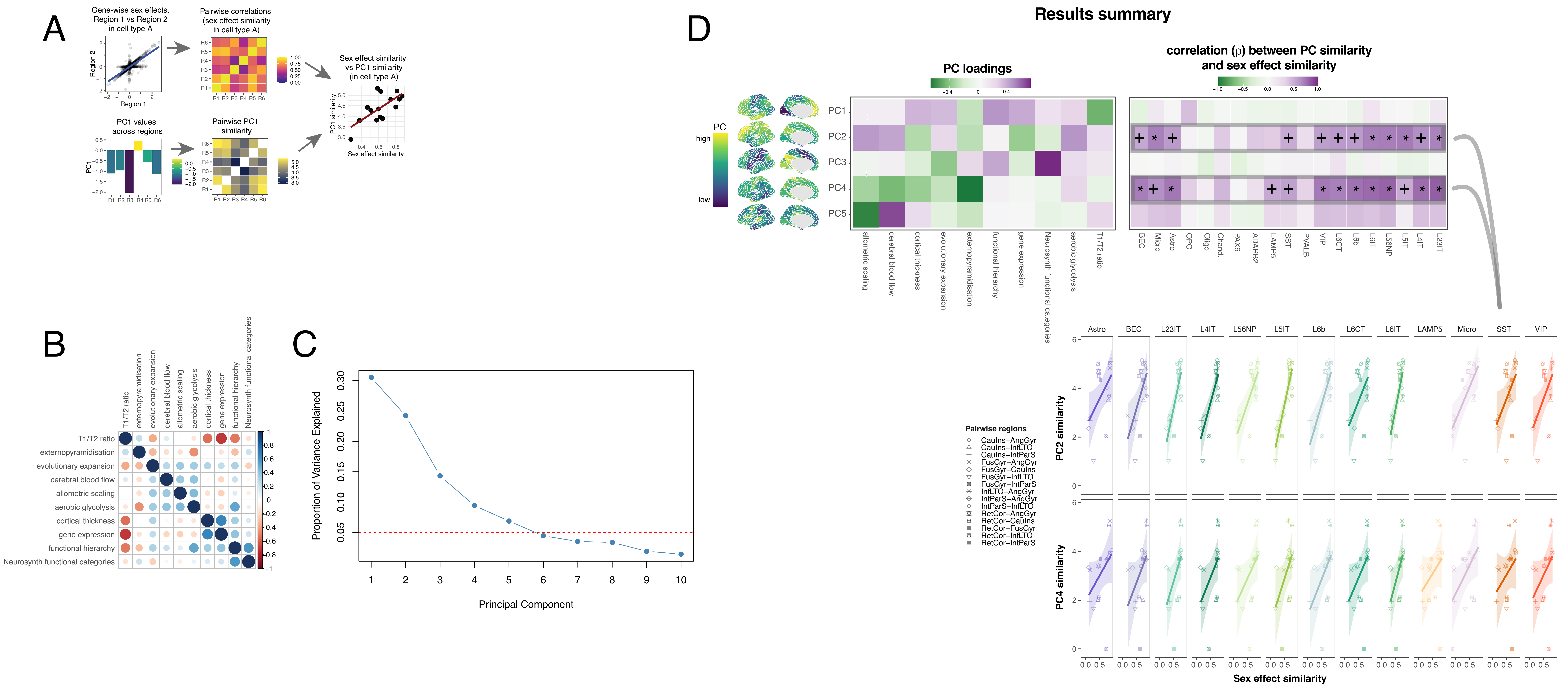
